# The role of MICOS in organizing mitochondrial cristae in malaria parasites

**DOI:** 10.1101/2025.10.13.682069

**Authors:** Silvia Tassan-Lugrezin, Irina Bregy, Judith López Orra, Nicholas I. Proellochs, Geert-Jan van Gemert, Rianne Stoter, Felix Evers, Taco W.A. Kooij, Laura van Niftrik

## Abstract

The malaria parasite mitochondria represent the only tractable, naturally occurring system in which cristae, the characteristic invaginations of the inner mitochondrial membrane, are known to be formed *de novo*. In traditional model organisms, the central complex involved in cristae organization is the mitochondrial contact site and cristae organizing system (MICOS). However, whether MICOS has an active role in the initial formation of cristae remains unclear. We identified two putative *Plasmodium falciparum* MICOS components, *Pf*MIC19 and *Pf*MIC60 and show that both genes are dispensable in asexual blood stages, gametocytes, and mosquito stages, albeit with a mild reduction in oocyst numbers. Immunofluorescence microscopy and electron tomography of gametocytes lacking either or both MICOS orthologues showed aberrant mitochondrial morphology and abnormal cristae, with a marked reduction of crista junctions. Thus, by utilizing the unique properties of *P. falciparum*, we confirmed the involvement of *Pf*MIC19 and *Pf*MIC60 in the organization of crista junctions, while providing evidence that MICOS is not required for the initial formation of cristae.

## Introduction

The mitochondrion is a double membrane-bound organelle found in most eukaryotes. Its primary function is to generate ATP through oxidative phosphorylation. The mitochondrion bears an outer mitochondrial membrane (OMM), which is a selectively permeable barrier to the cytoplasm, and an inner mitochondrial membrane (IMM), which separates the mitochondrial matrix from the intermembrane space. The IMM is divided into two distinct domains: the cristae membrane and the inner boundary membrane. The cristae membrane is highly convoluted and rich in ATP synthase and the electron transport chain complexes, thus forming a specialized area responsible for oxidative phosphorylation. The inner boundary membrane hosts proteins and protein complexes responsible for exchange of metabolites, protein translocation, and mitochondrial fusion (1-3). Both membrane regions are connected at the crista junctions. Crista junctions have been mainly described as tubular channels that connect the cristae membrane to the inner boundary membrane (4-6). Though, in organisms with lamellar cristae, such as yeast, crista junctions have also been identified as more elongated structures, also called slit-shaped crista junctions (7).

Crista junctions are known to be organized by a multi-subunit protein complex, the mitochondrial contact site and cristae organizing system (MICOS) (3, 8-11). Together with proteins of the OMM, MICOS is also part of the mitochondrial intermembrane space bridging complex that mediates contacts between the IMM and OMM (12-14). In humans, MICOS is organized into two protein sub-complexes integrated in the cristae membrane, the MIC10 and MIC60 sub-complexes, consisting of four and three different components, respectively (14-16). Disruption of either of the two MICOS sub-complexes has been shown to lead to loss of crista junctions and detachment of crista membranes. This generates aberrant cristae, followed by an atypical mitochondrial morphology (10, 15, 17). MIC60 is one of the key components of MICOS implicated in generating the negative curvature at the base of the cristae membrane (18, 19). MIC60 consists of an N-terminal transmembrane domain, responsible for anchoring it to the IMM, a coiled-coil domain and a mitofilin domain at the C-terminal end of the protein. These two C-terminal domains are essential to form and stabilize crista junctions through the tetramerization of MIC60 (19-21).

Strikingly, the mitochondrion of *Plasmodium falciparum*, the most virulent causing agent of human malaria, departs from the yeast or human model of mitochondria (22). Asexual blood-stage parasites (ABS) have a single acristate mitochondrion that harbours low levels of respiratory chain complex components and ATP synthase assemblies (23, 24). During development to the sexual, mosquito transmissible stages, also called gametocytes, the single mitochondrion separates into a set of 4-8 mitochondria (25) in which canonical mitochondrial processes, such as tricarboxylic acid cycle and oxidative phosphorylation gain relevance (23, 26-28). This is accompanied by a large increase in expression of ATP synthase and all relevant electron transport chain complexes but also by the *de novo* emergence of bulbous cristae (23, 25).

Even though mutation-induced aberrations of mitochondrial cristae morphology in *P. falciparum* have been sporadically reported (29), details on the molecular underpinnings of the bulbous shape and *de novo* biogenesis of cristae in *P. falciparum* gametocytes remain largely unknown. While a recent study in a related apicomplexan parasite, *Toxoplasma gondii*, has shown that the bulbous morphology is associated with highly divergent and large ATP synthase assemblies (5), the mechanisms involved in forming and maintaining stable crista junctions have yet to be investigated. Very little is known about any MICOS-like complex components or their potential roles in governing membrane architecture at the crista junction (29). Prior research has suggested PF3D7_1007800 as a putative orthologue of *MIC60* in *P. falciparum* (*PfMIC60*) based on sequence homology to the characteristic mitofilin domain, but until now, no other subunits of MICOS have been identified in *P. falciparum* (14). It is noteworthy that even though sequence homology and composition of MICOS diverges between distantly related eukaryotic clades, overall architecture is comparable to the opisthokont models (22).

Although MICOS has been described as an organizer of crista junctions, its role during the initial formation of nascent cristae has not been investigated. In this study, we used a combination of experimental genetics, high-resolution fluorescence microscopy, electron microscopy and electron tomography to elucidate the involvement of the parasite’s MICOS in cristae organization, with clear distinction from cristae formation.

## Results

### Identification of putative *Pf*MICOS components

To further investigate previously reported sequence homology of the MIC60 mitofilin domain to the *Pf*MIC60 candidate PF3D7_1007800 (14), we compared publicly available AlphaFold predictions of the protein sequence of the canonical MIC60 and our candidate sequence (Figure 1A, C and Figure S1A, C). In addition to the putative mitofilin domain in *Pf*MIC60, we identified a putative lipid binding domain and a coiled coil domain of similar length as described for opisthokonts MIC60. Interestingly, the annotated open reading frame of *Pf*MIC60 (UniProt Q8IJW5) is much larger than the *Saccharomyces cerevisiae* MICOS counterpart (UniProt A6ZZY0). The AlphaFold prediction of the full annotated open reading frame of PfMIC60 and ScMIC60 suggests the conserved domains (mitofilin, coiled coil and lipid binding domain) to make up the C-terminal half of PfMIC60. Even though confidence scores within the N-terminal half of the predicted molecule are low, Topology-based Evolutionary Domains (TED) identified a domain consisting of a stack of two parallel β-sheets in PfMIC60. Because this domain is predicted solely from computational analysis, both its actual existence in the native protein and its biological function remain unknown.

**Figure 1.**
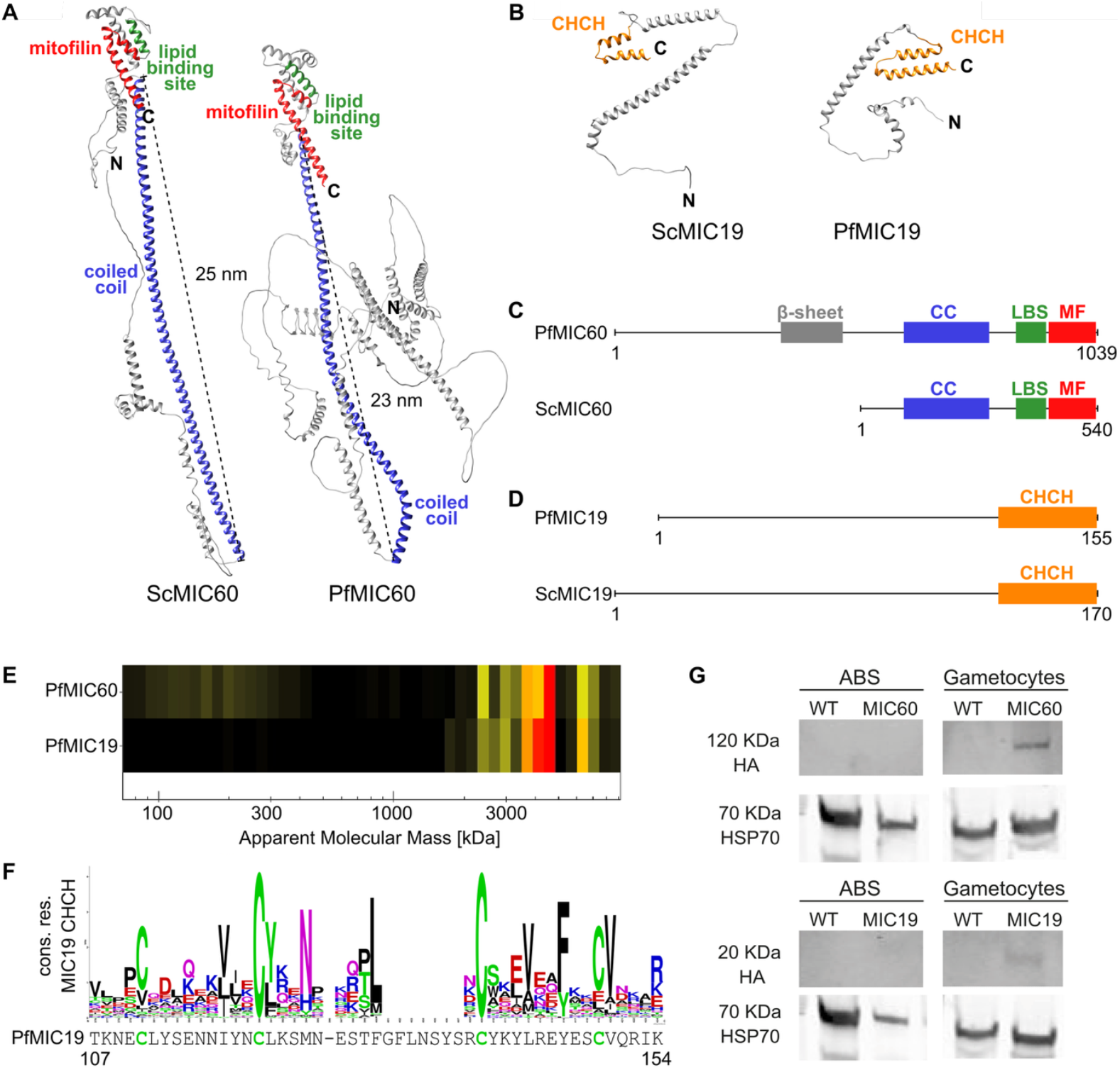
Identification of PfMICOS candidates based on predictive structural similarities, complexome profiling and the characteristic expression patterns expected for cristae modulators. A) AlphaFold predictions of ScMIC60 (AFDB ID: AF-A6ZZY0-F1-v4) and PfMIC60 (AFDB ID: AF-Q8IJW5-F1-v4)(32, 33). TED consensus domain predicted mitofilin domain (red), lipid binding site (green) and coiled coil domain (blue, predicted length annotated) are highlighted (34). N- and C-terminus are annotated. (local pLDDT confidence information in Figure S1). B) AlphaFold predictions of ScMIC19 (AFDB ID: AF-P43594-F1-v4) and PfMIC19 (AFDB ID: AF-Q8IIR5-F1-v4)(32, 33). TED predicted CC-helix-CC-helix (CHCH) domain is highlighted (34). N- and C-terminus are annotated. (local pLDDT confidence information in Figure S1). C) Schematic overview of the TED predicted domain architecture of ScMIC60 and PfMIC60. The additional domain annotated for PfMIC60 consists of ten antiparallel β-sheets of unknown function. D) Schematic overview of the TED predicted domain architecture of ScMIC19 and PfMIC19. E) Migration of PfMIC19/PfMIC60 across fractions of a 3-16% BN-PAGE gel. Corresponding apparent molecular mass of gel fractions is indicated on the x-axis. Relative abundance across fractions is indicated through a heatmap. Co-occurrence in gel fractions is indicative of a shared protein complex. F) Sequence logo derived form a MAFFT alignment of non-redundant protein sequences from MIC19 annotated entries listed on NCBI (MMseqs2 filtered at threshold 0.9, aligned sequences listed in Supplementary Information 2). Letter height indicates information content. Amino acid frequencies are shown by relative letter size. The characteristic set of two CX9C motifs found in MIC19 is conserved in PfMIC19 (sequence shown below). G) Western blot analysis of ABS and mature gametocytes of MIC19-HA and MIC60-HA. Tagged proteins were detected with rat anti-HA (PfMIC19 and PfMIC60, top) and rabbit anti-HSP70 was used as a loading control (bottom).

Since the *P. falciparum* genome lacks obvious orthologues of additional MICOS components, we re-examined our publicly available gametocyte-derived complexome profiling datasets (23). *Pf*MIC60 did not enter standard mass range gels but in samples separated on specialized, high molecular mass gels with lower acrylamide percentage, we found *PfMIC60* consistently comigrates with PF3D7_1108800 at an apparent mass of 4-5 MDa (Figure 1E, Supplementary Information 1) (23). PF3D7_1108800 is conserved across the *Plasmodium* genus (OG7_0323199) (30) and, while it lacks clear sequence homology outside of that, the AlphaFold prediction features a conserved CC-helix-CC-helix (CHCH) domain also found in MIC19, a known interactor of MIC60 in opisthokonts (Figure 1B, D and Figure S1B, C). Each of the two α-helices of the CHCH domain contains the conserved Cx9C motif that is characteristic for MIC19 CHCH domains in opisthokonts (Figure 1F). Based on these similarities and the comigration pattern, we propose to provisionally designate PF3D7_1108800 as *Pf*MIC19.

### *Pf*MIC60 and *Pf*MIC19 are expressed during gametocyte stages

To study the function of *Pf*MIC19 and *Pf*MIC60 as possible MICOS components, we analysed their expression profile and localization in ABS and gametocytes. Therefore, we generated two transgenic parasite lines, *mic19-HA* and *mic60-HA* in which we introduced a *3xHA-GlmS* tag at the 3’ end of the respective genes. The transgenic lines also contain a mitochondrially targeted mScarlet for visualization of the organelle and protein co-localization (Figure S2A). Correct integration and absence of wild-type (WT) parasites was confirmed through diagnostic PCR (Figure S2B). Western blot analysis of mixed ABS and mature gametocyte samples revealed that *Pf*MIC19 (19.7 kDa) and *Pf*MIC60 (120 kDa) are undetectable in acristate ABS but expressed in the cristate sexual blood stages (Figure 1G). As a positive control for mitochondrial localization, we used the MitoRed line (31), which expresses the mitochondrially targeted mScarlet from a silent intergenic locus. Unfortunately, attempts to localise *Pf*MIC19-HA and *Pf*MIC60-HA using conventional immunofluorescence microscopy and ultra-expansion microscopy were unsuccessful, likely due to their naturally low abundance (Figure S2C-D) highlighted by our western blot analysis (Figure 1G).

### *Pf*MIC19 and *Pf*MIC60 are not essential for *P.falciparum in vitro* life-cycle progression

To investigate the essentiality of *Pf*MIC19 and *Pf*MIC60 during the *P. falciparum* life cycle, we generated the knockout parasite lines *mic19*^*–*^ and *mic60*^*–*^ in which the respective genes were replaced by a cytoplasmic green fluorescent protein (GFP) (Figure S2A). In addition, we generated the *mic19/60*^*–*^ double-knockout line in which *PfMIC60* was replaced by a cytoplasmic red fluorescent protein (mCherry) in the *mic19*^*–*^ background (Figure S3A). Diagnostic PCR confirmed proper integration and absence of WT parasites in all the generated lines (Figure S2B and Figure S3B). To assess parasite viability, we performed a standard growth assay on ABS. Parasite replication was followed for two replication cycles, revealing that ABS of *mic19*^*–*^, *mic60*^*–*^, and *mic19/60*^*–*^ grow at a comparable rate to parental parasites (NF54) (Figure 2A). Interestingly, *mic19*^*–*^, *mic60*^*–*^, and *mic19/60*^*–*^ also produced mature gametocytes with no apparent delay. To verify their viability, we performed exflagellation assays that revealed no difference in maturation compared to the NF54 parental control (Figure 2B). In addition, gametocytes were fed to mosquitoes through standard membrane feeding assays. All lines infected mosquitoes and we observed a reduction in oocyst numbers in *mic19*^*–*^, *mic60*^*–*^, and *mic19/60*^*–*^ compared to NF54 (Figure 2C and Figure 2D). The high variability encountered in the standard membrane feeding assays, though, partially obstructs a clear conclusion on the biological relevance of the observed reduction in oocyst numbers. Nevertheless, sporozoite production was confirmed for all generated knockout lines, indicating that *Pf*MIC19 and *Pf*MIC60 are not essential for the parasites to complete mosquito-stage development (Figure S5). In summary, under *in vitro* culture conditions, *Pf*MIC19 and *Pf*MIC60 are dispensable for survival and development of ABS and gametocytes. Absence of these MICOS components is also compatible with progression through mosquito-stage development.

**Figure 2.**
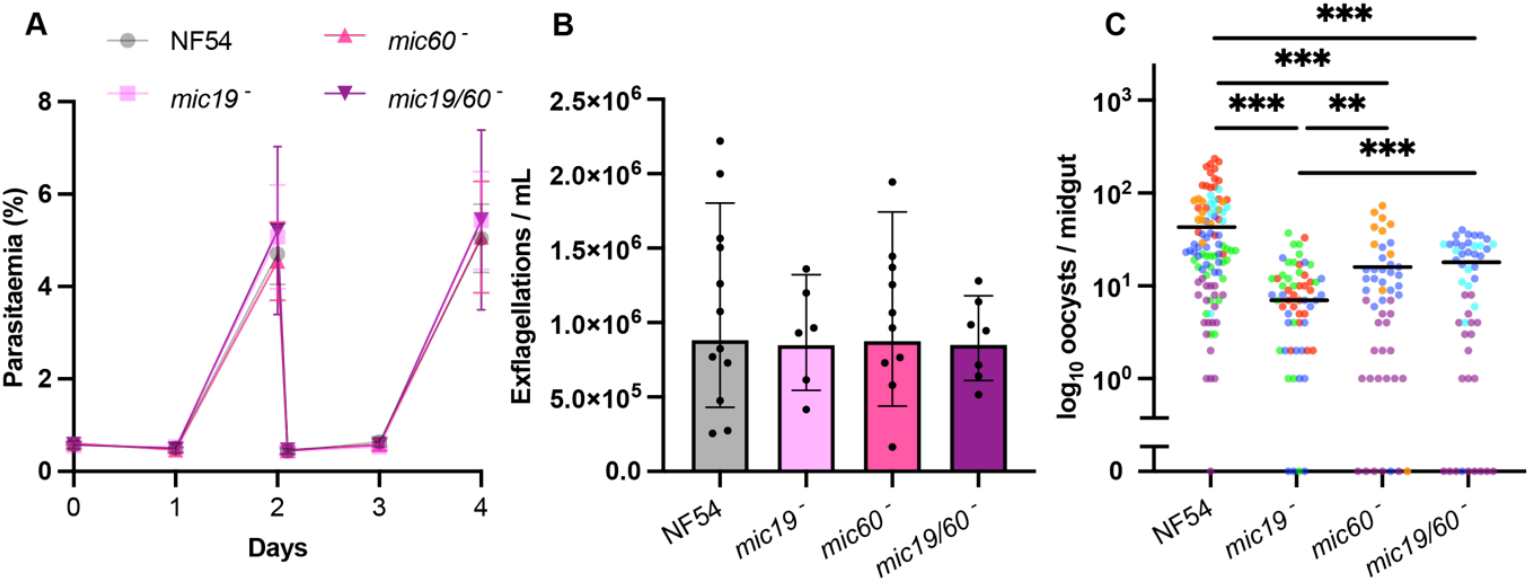
Consequences of deletion of PfMIC60 and PfMIC19 on differentiation and survival of P. falciparum. A) Growth curve of NF54, mic19^−^, mic60^−^, and mic19/60^−^ ABS over time (in days). (n = 3 for NF54, mic19^−^, mic60^−^ and n = 2 for mic19/60^−^). B) Exflagellation assay of NF54, mic19^−^, mic60^−^, and mic19/60^−^. (n >= 6). Statistical significance calculated with one-way ANOVA with Sidàk correction for full comparison between all cell lines. C) Infection of A. stephensi mosquitoes with NF54, mic19^−^, mic60^−^, and mic19/60^−^ in 6 independent experiments indicated in different colours. Means (black line): NF54 = 43, mic19^−^ = 7.0, mic60^−^ = 15.0 and mic19/60^−^ = 17.1. Statistical significance was assessed with Wald statistics and indicated as follows: ^*^p < 0.05, ^**^p < 0.01, ^***^p < 0.001, ^****^p < 0.0001. Oocyst count of 0 was plotted at the x-axis for visualisation purposes.

### Knockout of *PfMIC60* and *PfMIC19* impairs mitochondrial morphology in gametocytes

Next, we investigated the impact of *Pf*MIC19 and *Pf*MIC60 absence on mitochondrial morphology. Due to the lack of detectable protein expression in ABS, we focused on mature gametocytes, where cristae are abundant and protein expression was detected in western blot. We stained mitochondria of parental NF54 line, single and double knockout line mature gametocytes with MitoTracker and used alpha tubulin II as a known and previously confirmed male marker (35). To analyse the variability of normal mitochondrial morphology in the NF54 background, we quantified the imaging data using the Mitochondria-Analyzer plug-in for Fiji (36, 37). MitoTracker signals were classified into three categories: (1) tubular, when only one/two elongated and highly branched mitochondrial signals were identified per cell body; (2) intermediate, when multiple partially branched mitochondrial signals were observed; and (3) rounded-up, when mitochondrial signals appeared smaller, more spherical, and dispersed (Figure 3A). Note that light microscopy does not resolve individual mitochondria if they are in close contact with each other (25). Nevertheless, the analysis revealed that mitochondrial morphology of both male and female gametocytes was partially altered in all knockout lines, characterized by a predominance of rounded-up and intermediate mitochondria, and a marked reduction of tubular networks. (Figure 3B, Figure S6A). In addition to the rounded-up appearance of mitochondria, we also observed a dispersion of the mitochondrial network into spatially separated clusters. Wild-type cells contain a single compact cluster of closely apposed tubular mitochondria, whereas knockout cells display multiple smaller, dispersed mitochondrial clusters. Based on the number of individually identifiable MitoTracker signals within a cell, we determined the relative dispersion of the mitochondrial network. Even though not significant, the data indicates a trend towards increased network dispersion in all knockout lines (24%, 21%, and 23% in *mic19*^−^, *mic60*^−^, and *mic19/60*^−^) (Figure S6B). The single identified mitochondrial signals were ∼30% smaller (Figure S6C), more spherical (36%, 24%, and 20%) (Figure 3C), and less branched (50%, 55%, and 40%) (Figure S6D) compared to the parental NF54 line. In addition, total mitochondrial volume per cell was reduced in all knockout lines with 15%, 17%, and 13%, respectively (Figure 3C). Interestingly, deletion of both genes did not exacerbate the mitochondrial phenotype compared to the single knockouts. Finally, no difference was detected in MitoTracker signal intensity, possibly indicating that the membrane potential is not affected in the single nor double knockouts (Figure S6E). In conclusion, while NF54 cells contain mainly tubular mitochondria, *mic19*^−^, *mic60*^−^, and *mic19/60*^*–*^ gametocytes have more intermediate and rounded-up mitochondria.

**Figure 3.**
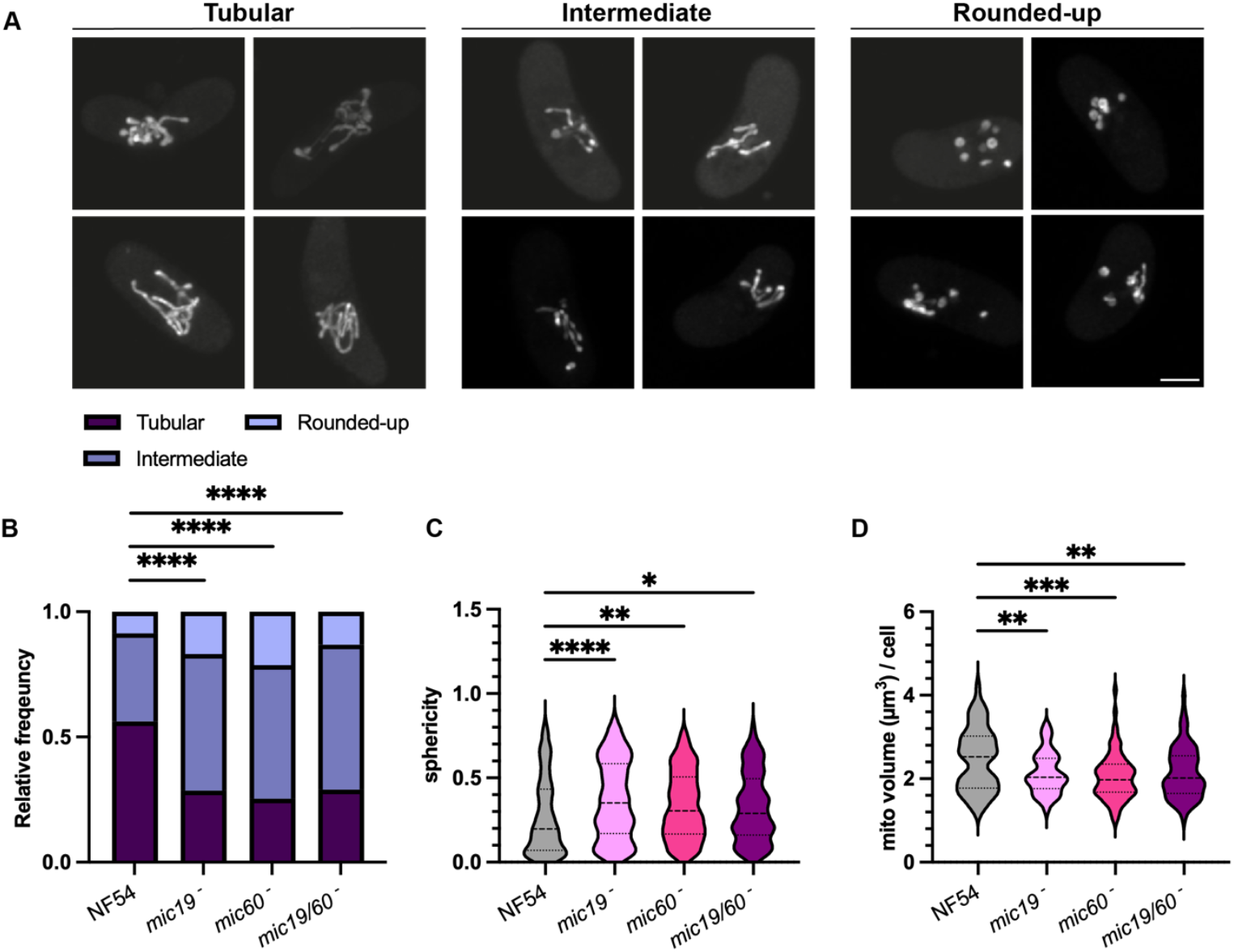
Fluorescence microscopy reveals aberrant mitochondrial morphology in mature P. falciparum mic19^−^, mic60^−^, and mic19/60^−^ gametocytes. A) Representative fluorescent microscopy images of mature NF54 gametocytes with tubular, intermediate, or rounded-up mitochondria. Samples were stained with MitoTracker. Images are maximum intensity projections of Z-stack confocal Airyscan images. Scale bars: 2 µm. B) Analysis of mitochondrial shape based on mean branch length per cell. Rounded-up = branch length < 2 μm, intermediate = branch length between 2 μm and 10 μm, tubular = branch length > 10 μm. Statistical analysis using Chi-squared test with Bonferroni correction for full comparison between all cell lines; significance is indicated as follows: ^*^p < 0.05, ^**^p < 0.01, ^***^p < 0.001, ^****^p < 0.0001. C) sphericity of mitochondria, D) mitochondrial volume per cell. In all cases n° of replicates > 3 with total number of cells ≥ 75. Statistical significance calculated with one-way ANOVA with Sidàk correction for full comparison between all cell lines; significance is indicated as follows: ^*^p < 0.05, ^**^p < 0.01, ^***^p < 0.001, ^****^p < 0.0001.

### *Pf*MIC60 and *Pf*MIC19 are involved in shaping mitochondrial cristae

To assess the role of *Pf*MIC19 and *Pf*MIC60 in cristae morphology, we performed transmission electron microscopy (TEM) on mature gametocyte of *mic19*^−^, *mic60*^−^, *mic19/60*^−^, and the parental NF54. Two-dimensional analysis of thin sections revealed cristae deformations in all knockout lines (Figure 4A). We did not observe differences in cristae morphology between male and female parasites (Figure 4A and Supplementary Information 2). Length, diameter, and area occupied by individual cristae from over 30 mitochondrial cross-sections per sample were measured (Figure 4B-D). Note that such measurements do not indicate the true total length or diameter of cristae, as the data is two-dimensional. The recorded values are to be considered indicative of trends, rather than absolute dimensions of cristae. Even though cristae of the two single knockout lines (*mic19*^−^ and *mic60*^−^) appeared disorganized, significant increase in apparent length (but not diameter) of cristae was only detected in *mic19/60*^−^ gametocytes (with a mean crista length of 155 nm compared to 96 nm for NF54) (Figure 4B-C). Despite the lack of overall significance of these changes in the single knockouts, the spread of recorded crista length did increase in all knockout lines, therefore indicating a trend towards heterogeneity of crista architecture. In line with those observations, crista area increased in all knockout lines (Figure 4D). We expect this effect to translate into the third dimension and thus conclude that the mean crista volume increases with the loss of either *PfMIC19, PfMIC60*, or both. Even though the deletion of *PfMIC19* and/or *PfMIC60* clearly affected crista morphology, mean cristae coverage per mitochondrial cross-section did not significantly change in *mic19*^−^ and *mic60*^−^ (Figure 4E). The double knockout line (*mic19/60*^−^), however, reached significance due to a small population of severely aberrant cristae (Figure 4F and Figure S7). We observed instances of extremely long cristae, as well as mitochondria with large single- or double-membraned circles within the mitochondrial cross-section. The reason for the lack of a significant change in *mic19*^−^ and *mic60*^−^ despite the increased area per crista can be calculated from the same datapoints and shows that the mean number of crista cross-sections per µm^2^ of mitochondrial cross-section drops from 33.8 in NF54 to 20.3, 16.3 and 26.4 in *mic19*^−^, *mic60*^−^ and *mic19/60*^−^, respectively. These effects in combination suggest that the increased area per crista is associated with a reduced number of cristae per mitochondrial area. Given the observed aberrations in crista morphology in knockout lines, we conclude that membrane invaginations can still form, even in the absence of *Pf*MIC19 and *Pf*MIC60, suggesting that these proteins are not essential for initiating crista formation. Nevertheless, *Pf*MIC60 and *Pf*MIC19 both play an essential role in shaping nascent cristae. Furthermore, our data indicates the effect size increases with simultaneous ablation of both proteins.

**Figure 4.**
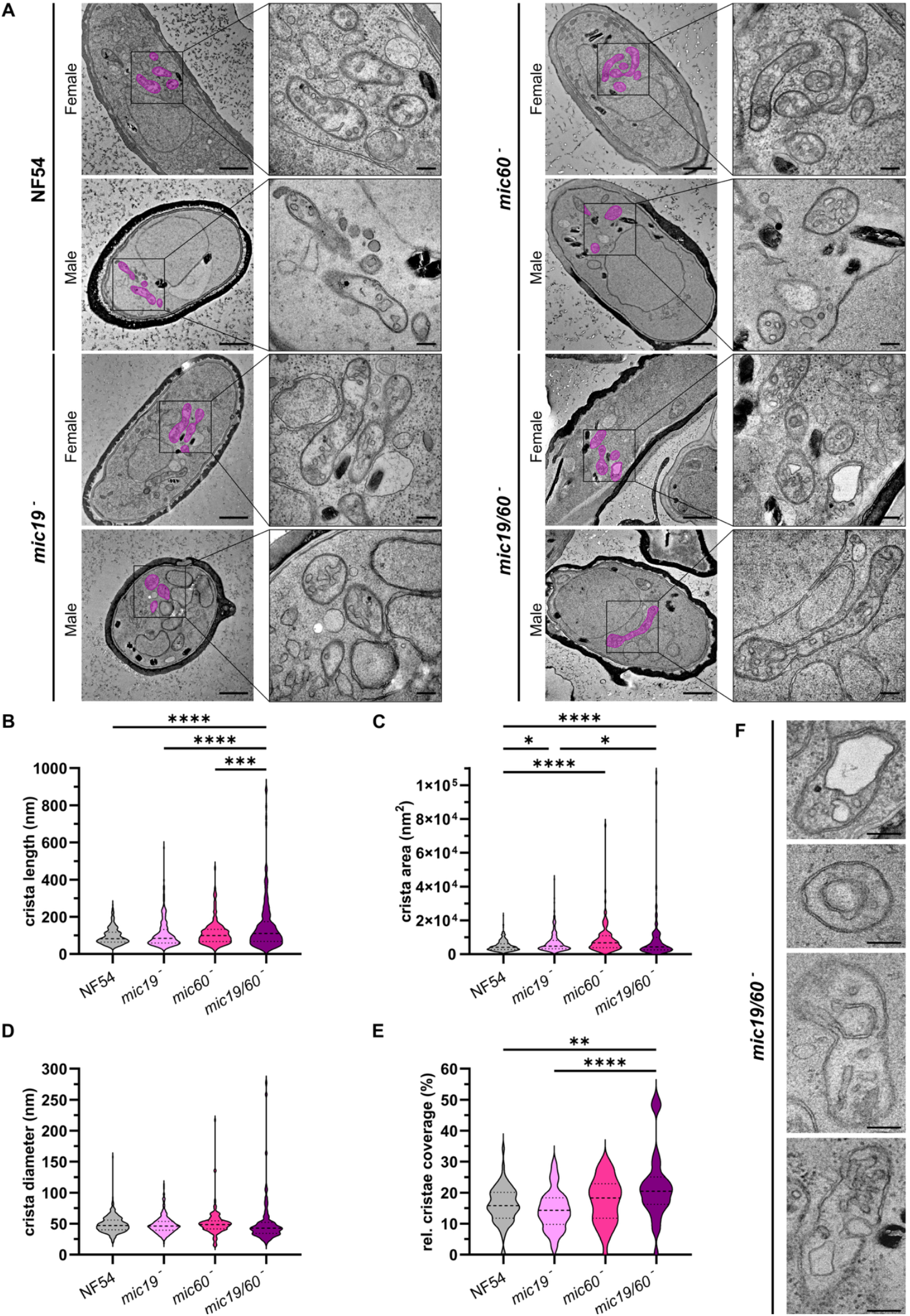
Transmission electron microscopy reveals aberrant crista structures in mature P. falciparum mic19^−^, mic60^−^, and mic19/60^−^ gametocytes. A) Representative micrographs of NF54 (WT), mic19^−^, mic60^−^, and mic19/60^−^ gametocytes. Mitochondria within a rectangular selection are highlighted in pink. Zoom-ins on these selections are shown on the right side of each micrograph. For each cell line, a male and a female parasite are shown. Scale bars: 1 µm / 200 nm. B - D) Quantification of the length (log) (B), width (log) (C), and area (log) (D) of cross**-**sections through individual cristae (n ≥ 100, from ≥ 30 mitochondria). E) Quantification of cristae coverage based on the total crista area of a mitochondrial cross section relative to the area of the respective mitochondrial cross section (n ≥ 30). F) Examples of severely aberrant mitochondria as observed in TEM of mic19/60^−^ gametocytes. Scale bars: 200 nm. Statistical analysis of B – E: one-way ANOVA with Sidàk correction for full comparison between all cell lines; significance is indicated as follows: ^*^p < 0.05, ^**^p < 0.01, ^***^p < 0.001, ^****^p < 0.0001.

### *Pf*MIC60 and *Pf*MIC19 are important for proper crista junction formation

To delve deeper into the effects of *PfMIC19* and *PfMIC60* deletions, we performed dual-tilt TEM tomography on 250 nm sections of the same samples used in Figure 4. The phenotypes matched our observations from two-dimensional acquisitions. We found crista morphology to be altered in all knockout lines, with most severe aberrations found in *mic19/60*^−^ (Figure 5A-B). We manually segmented one representative mitochondrion per parasite line to visualize the observed effects (Figure 5C and Movies 1-4). We observed that in all knockout parasites, we rarely identified connections between the crista membranes and the boundary membrane. To quantify the observed loss of membrane connectivity, we analysed the connectivity of all cristae within mitochondrial volumes from different gametocytes (Figure 5E-F). We thoroughly searched for crista junctions of each crista found within the mitochondrial segments. Note that these mitochondrial volumes are not full mitochondria, but large segments thereof. As a result of the incompleteness of the mitochondria within the section, and the tomography specific artefact of the missing wedge, we were unable to confirm whether cristae were in fact fully detached from the boundary membrane, or just too long to fit within the observable z-range. Instead, we quantified the fraction of cristae for which we identified a crista junction (Figure 5E). While we identified crista junctions for 55% of the cristae seen in NF54 gametocytes, we could only confirm crista junction presence 9% of the cristae traced in *mic19*^−^ and 23% of the cristae in the *mic60*^−^ sample. Similarly, connectivity also dropped in the double knockout, with 21% of confirmed crista junctions. Note that for *mic19/60*^−^ we chose to measure connectivity in mitochondria that we considered representative for the population average rather than the extreme cases shown in (Figure 4F and Figure S7). For each parasite line, we measured the diameter of at least 12 crista junctions (Figure 5E-F). A noteworthy, yet non-significant drop of the crista junction diameter was observed in *mic19/60*^−^, where the mean crista junction diameter dropped from 25 nm in the NF54 control to 20 nm in *mic19/60*^−^. Collectively, these results demonstrate that the deletion of *PfMIC19* and/or *PfMIC60* leads to a reduced number of crista junctions and a possible reduction in crista junction diameter in *mic19/60*^−^.

**Figure 5.**
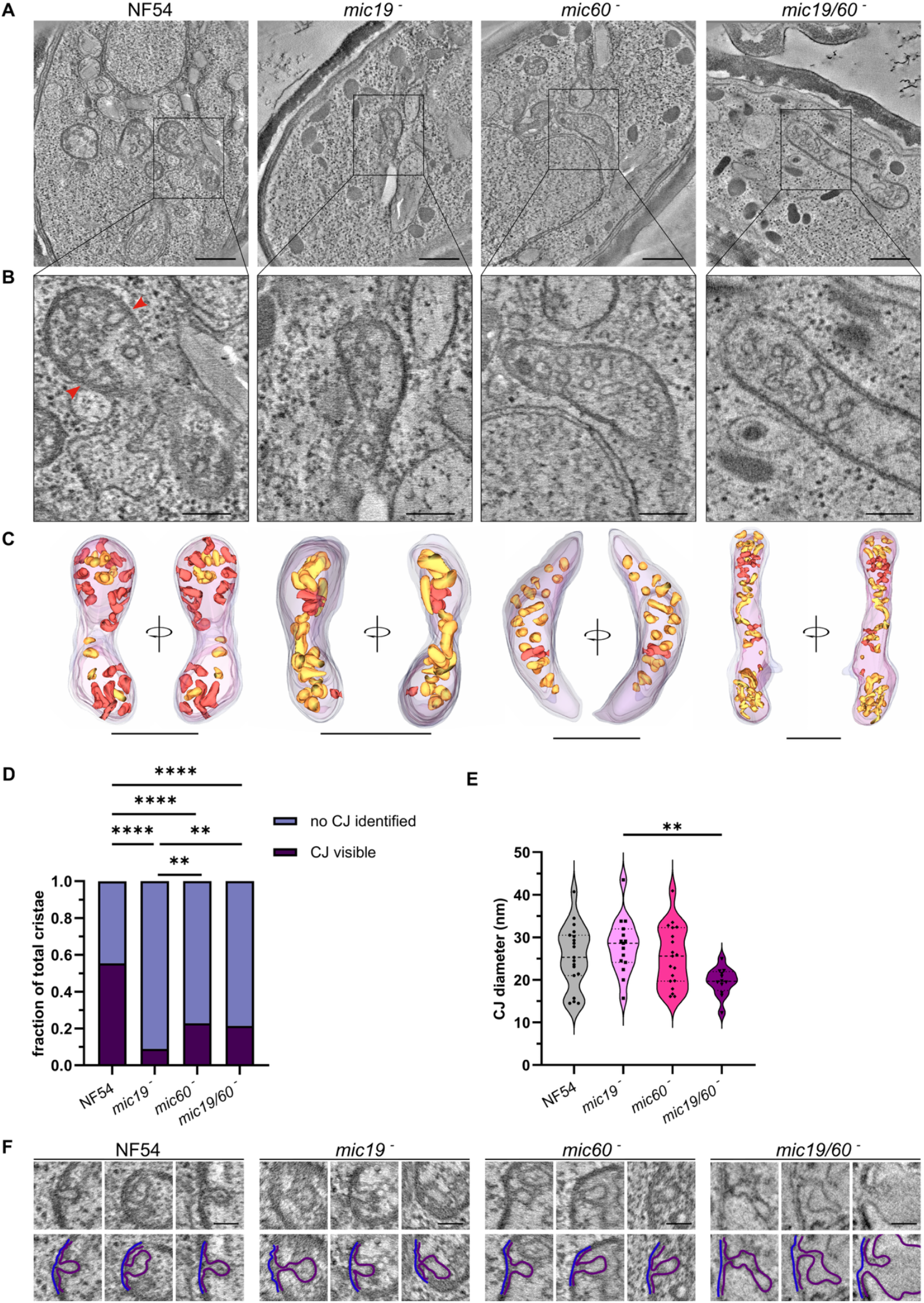
Transmission electron tomography reveals a drop in crista junction prevalence in mature P. falciparum mic19^−^, mic60^−^, and mic19/60^−^ gametocytes. A) Representative snapshots through TEM tomograms of mature NF54 (WT), mic19^−^, mic60^−^, and mic19/mic60^−^ gametocytes. Scale bars: 500 nm. B) Zoom-ins of individual mitochondria from the tomograms shown in A. Crista junctions highlighted with red arrow heads. Scale bars: 200 nm. C) 3D model of 250 nm mitochondrial sections of the mitochondria shown in B. Each model is shown from two angles 180° to each other. Scale bars: 500 nm (full tomograms and mitochondrial segmentations in Movies 1-4). For red coloured cristae, a CJ was identified within the inspected 250 nm section of the mitochondrion. For yellow-coloured cristae we could not identify connection to the boundary membrane within the tomogram. D) Quantification of the relative proportions of cristae for which a crista junction (CJ) was observed within the 250 nm section (n ≥ 56). Statistical analysis using Chi-squared test with Bonferroni correction for full comparison between all cell lines; significance is indicated as follows: ^*^p < 0.05, ^**^p < 0.01, ^***^p < 0.001, ^****^p < 0.0001. E) Quantification of the diameter of crista junctions at their narrowest point (n ≥ 12, from several mitochondria per sample). Statistical analysis using one-way ANOVA with Sidàk correction for full comparison between all cell lines; significance is indicated as follows: ^*^p < 0.05, ^**^p < 0.01, ^***^p < 0.001, ^****^p < 0.0001. F) Three representative crista junctions from mature NF54, mic19^−^, mic60^−^, and mic19/mic60^−^ gametocytes. Scale bars: 100 nm. 2D segmentation of the outer mitochondrial membrane (blue) and the inner mitochondrial membrane (violet) shown in the bottom row.

## Discussion

MICOS is a protein complex of the inner mitochondrial membrane that is involved in the organization of mitochondrial cristae and stabilization of crista junctions (3, 8-10). Though well established as the cristae organizing system in opisthokonts (the clade including animals and fungi), MICOS has not previously been studied in apicomplexan parasites (38). The human malaria parasite *P. falciparum*, offers a novel perspective on cristae biology, as it is one of very few species that reportedly grow cristae *de novo* from an acristate mitochondrion (25, 39, 40). Due to its medical relevance and the central role of the mitochondria as a validated drug target (41), *P. falciparum* is an invaluable model to study the role of MICOS in cristae organization and formation. Recent bioinformatic data revealed a putative *Pf*MIC60, thus priming the investigation of other components of the complex (14, 23). In this study, we investigated the roles of the putative *PfMIC60* and *PfMIC19* orthologues in the formation and organization of cristae.

The *P. falciparum* mitochondrial proteome intensely varies between different life cycle stages. In fact, almost all proteins involved in oxidative phosphorylation are dramatically increased in gametocyte stages compared to ABS (40, 42). The same trend was observed for *Pf*MIC19 and *Pf*MIC60. Although transcriptomic data suggested that low-level transcripts of *PfMIC19* and *PfMIC60* might be present in ABS (43, 44), we did not detect either of the proteins in ABS by western blot analysis. The apparent absence could indicate that transcripts are not translated in ABS or that the proteins’ expression was below detection limits of western blot analysis. In contrast, both proteins were detectable, albeit in low abundance, in mature gametocytes. The expression patterns of *Pf*MIC19 and *Pf*MIC60 are consistent with the absence of cristae in ABS and their appearance during gametocytogenesis (25, 40). The low abundance of *Pf*MICOS components even in mature gametocytes likely contributed to our difficulties confirming mitochondrial localization of *Pf*MIC19 and *Pf*MIC60 using fluorescence microscopy.

The apparent absence of *Pf*MIC19 and *Pf*MIC60 in ABS also explains the lack of a growth defect in this part of the life cycle. Surprisingly though, gametocytes and mosquito stages, that are believed to strongly rely on oxidative phosphorylation for their survival (40, 42), were largely unaffected by depletion of these MICOS components, with the exception of a slight reduction in mosquito colonization. This lack of growth arrest has also been described for the knockout of some MICOS components in human cells, yeast and *Trypanosoma brucei* (16, 45-47). Though, in some cases, MICOS subunits were shown to be essential for cell survival and growth, such as the cryptic mitofilin MIC34 and MIC40 in *T. brucei* (48). In addition, disruption of some MICOS subunits has been reported to partially impair cell respiration (10, 20, 49, 50). Considering these relationships, successful propagation of mosquito stages suggests either that the growth defect is masked by the unnaturally rich environment encountered in blood culture medium (51) or that the reported impact on respiratory complexes does not occur to a relevant extent in *P. falciparum* which is supported by the absence of detectable differences in membrane potential .

Consistent with prior observations in human cells (10, 52), knockouts of *Pf*MIC19 and *Pf*MIC60 strongly altered mitochondrial morphology. In fact, both single and the double knockout parasite lines exhibited mitochondria that were more rounded-up and spherical. The presence of rounded-up mitochondria upon knockout of MIC60 and MIC19 has also been observed in human and yeast cells and appointed as a physiological response to mitochondrial stress (10, 20). Given this background and the extreme variability in mitochondrial shape we assume mitochondrial rounding-up in *P. falciparum* is also a consequence of mitochondrial stress, due to the loss of *Pf*MIC19 and *Pf*MIC60, rather than a direct involvement of these proteins in mitochondrial shape. Our observation of this mitochondrial rounding-up in the NF54 parental control, could indicate a baseline level of mitochondrial stress in culture conditions. Similarly, we cannot clearly appoint the reduction in oocyst number to a direct response to the loss of *Pf*MIC19 and/or *Pf*MIC60, as it may reflect mitochondrial stress or even result from the high variability enclosed in the assay itself. Still, it is striking that, despite the pronounced morphological phenotype, and the possibly high mitochondrial stress levels, the parasites appeared mostly unaffected in life cycle propagation, raising questions about the functional relevance of crista architecture and overall mitochondrial morphology at these stages.

The MitochondriaAnalyzer plug-in for FIJI, used here for the first time on *P. falciparum* samples, proved to be a potent tool to improve bulk quantification of fluorescence microscopy data (36). Of course, it is also important to consider that while fluorescent microscopy is a powerful technique, it cannot reliably determine the number of mitochondria in *P. falciparum* gametocytes. As shown by TEM data, mitochondria are often located in very close proximity (25, 53), thus the fluorescence-based mitochondrial counts reflect more on whether mitochondria are sufficiently dispersed to be recognized as separate, rather than providing an absolute count.

The depletion of either or both *P. falciparum* MICOS genes leads to alterations in the organization of cristae membranes while total cristae coverage remains largely unaffected, thus suggesting that membrane invaginations still arise but are not properly arranged in these knockout lines. While the extent of cristae disruptions in *mic19*^*–*^ and *mic60*^*–*^ gametocytes was comparable (as shown in thin section TEM and TEM tomography), the double knockout parasites showed stronger morphological defects compared to the single knockouts. This may suggest that in *P. falciparum* the depletion of just one of these proteins does not fully disrupt MICOS function, Instead, our data indicates progressive loss of MICOS function upon the depletion of multiple MICOS components.

While MIC60 knockout in human cells showed an almost complete loss of crista junctions (10), cristae aberrations were more subtle in our knockout lines. Nevertheless, we did see a decrease in crista junction abundance in all lines, and a trend that is indicative of a reduction of crista junction diameter in the double knockout. Interestingly, in both human and yeast cells, crista junctions exhibit diameters ranging from 12 to 40 nm, with an average of 28 nm, which is consistent and comparable with our observation in the parental NF54 gametocytes (54-56). The change in crista junction diameter suggests that, at least in *mic19/60*^*–*^, the remaining crista junctions are affected by the loss of *Pf*MIC60 and *Pf*MIC19. It remains uncertain, though possible, that protein-mediated stabilization is entirely absent in the double knockout.

It is important to note that effects on crista junction structure upon destabilization of the MIC60 subcomplex is a phenomenon described also in yeast and human cells. (10, 19). Previous studies showed that depletion of *MIC60* in HeLa cells led to an increase in crista junction diameter, while the depletion of *MIC19* did not affect this parameter (11). In contrast to that, destabilization of the MIC60 tetramer in yeast led to a decrease in crista junction diameter (19). Interestingly, our findings do not suggest effects on crista junction diameter for either one of the single knockouts, while the double knockout resulted in a 20% reduction. The central, tetrameric coiled coil domain of MIC60 has been suggested to define the diameter of the crista junction (19). While not experimentally proven, MICOS induced stabilization of crista junction diameter is a plausible mechanism also for *P. falciparum*. In fact, the mean diameter of *P. falciparum* crista junctions aligns well with the theoretical length of the coiled coil domain predicted by AlphaFold.

Collectively, we have demonstrated that *Pf*MIC19 and *Pf*MIC60 are putative mitochondrial proteins expressed in gametocytes. Their deletion does not affect ABS nor sexual blood stages in standard culture conditions. While oocyst counts were modestly reduced, sporozoite production did not appear to be affected, indicating that the proteins are not essential at any point of the parasite life cycle investigated in this study. Still, knockout of *Pf*MIC19 and *Pf*MIC60 strongly altered mitochondrial morphology in gametocytes. Mitochondrial cristae are aberrant but present, with a reduction in crista junction abundance, confirming a role in *Pf*MICOS. Although molecular mechanisms of membrane bending and anchoring are yet to be elucidated, our study demonstrated the existence of *Pf*MICOS. In contrast to other systems where MICOS has been described, *P. falciparum* further allows us to clearly decipher cause-effect relationships in this context. We conclude that even in the absence of *Pf*MIC60 and *Pf*MIC19, cristae are formed *de novo* and allowing for parasite survival. Thus, our data provide direct evidence that components of *Pf*MICOS are essential for crista junction organization and stabilization, but not for initial cristae formation. This functional separation is likely to apply beyond *P. falciparum*, to evolutionary distant organisms such as human and yeast (22). Our results highlight the unique utility that the two-state mitochondrial architecture in *P. falciparum* provides to the field of mitochondrial biology and lays the groundwork for future studies on the causal dissection of crista biogenesis during *P. falciparum* gametocytogenesis and beyond.

## Material and Methods

### Plasmid generation

To generate the *mic19-HA* and *mic60-HA* homology-direct repair (HDR) plasmids, the pRF0139 empty tagging plasmid was used, containing 3xHA-GlmS, PBANKA_142660 bidirecoonal 3’UTR, and the mitochondrial-mScarlet marker (31). 5’ and 3’ homology arms (HRs) were amplified from genomic NF54 DNA and cloned with specific restricoon enzymes (Table S1).

HDR templates for *mic19*^−^ and *mic60*^−^ were generated from pGK backbone (57), which contains a pBAT backbone (58) with H2B promotor, GFP and PBANKA_142660 bidirecoonal 3’UTR. The 5’ and 3’ HRs, amplified from NF54 gDNA, were cloned into pGK using specific restricoon sites (Table S1).

To generate the HDR template for the *mic19/60*^−^, 5’ and 3’ HR were inserted into pCK2a, which contains a pBAT backbone (58) with H2B promotor, mCherry and PBANKA_142660 bidirecoonal 3’UTR.

Cas9 guide plasmids were generated inserong a pair of annealed complementary, guide encoding, oligonucleoodes into pBF-gC (59) (kind giq from Till Voss) or, pRF0568, a plasmid derived from pMLB626 (60) (kind giq from Marcus Lee) without a 3xHA tag fused to *Cas9*. BsaI and BbsI, respecovely, restricoon enzymes were used to insert the oligonucleoodes and generate the final guide plasmids (Table S1). All PCR reacoons were performed using Primestar GXL DNA polymerase (Takara Bio), and all cloning enzymes were provided by New England Biolabs.

### *P. falciparum* culturing and transfection

*P. falciparum* ABS were cultured in RPMI medium (with 25 mM HEPES, 100 mM hypoxanthine, and 24 mM sodium bicarbonate) completed with 10% AB+ human serum and 5% 0+ human Red Blood Cells (RBCs) (Sanquin, The Netherlands) (61). The cultures were kept at 37° C in a low oxygen environment (3% O_2_, 4% CO_2_). When needed, ring-stage parasites synchronization was achieved using sorbitol treatment (62). Culture parasitaemia, was monitored with Giemsa-stained blood smears using a 100X oil objective.

For transfections 60 µg of each Cas9 guides and linearized repair plasmids were ethanol precipitated and transfected on a ring-stage synchronized culture, as previously described (63). Briefly, the plasmids were dissolved and mixed with cytomix (0.895% KCl, 0.0017% CaCl_2_, 0.076% EGTA, 0.102% MgCl_2_, 0.0871% K_2_HPO_4_, 0.068% KH_2_PO_4_, 0.708% HEPES) for a total volume of 300 µl. The DNA was then added to 0.4 ml of packed ring-stage infected RBCs, and the solution was electroporated (0.31 kV and 950 μF). Parasites were left to recover for 4 h after transfection in complete medium and after that treated in 2.5 nM WR99210 (Jacobus Pharmaceuticals) or 40 µg/ml blasticidin (Gibco #R210-01) for 5 days. To obtain an isogenic population, parasites were isolated using flow cytometry (64) based on GFP (*mic19*^−^ and *mic60*^−^), mCherry (*mic19/60*^−^) or mScarlet (*mic19-HA* and *mic60-HA*) expression, using a CytoFLEX SRT benchtop Cell Sorter (Beckman Coulter). Transfected parasites were cultured until recovered and integration was confirmed by diagnostic PCR (Table S1).

### Growth assay

Parasite cultures were tightly synchronized through a double sorbitol synchronisation performed at 0 h and 16 h. Ring-stage parasites were determined through flow cytometry and cultures were diluted to have a 0.5% final parasitaemia. Cultures were set up at 0.5% parasitaemia and samples were harvested for 4 consecutive days. Samples for flow cytometry were taken directly after set-up, on day 1, 2 (before and after dilution), 3 and 4. On day 2 all cultures were diluted with the same dilution factor (1/20). When harvested, samples were fixed in 0.25% glutaraldehyde (ITW Reagents, CA0589.100) in PBS and, before flow cytometry (CytoFLEX - 13-color, Beckman Coulter), stained with SYBR™ green (ThermoFisher, 7563) 1:10.000 for 30 min at 37° C. Data was analysed using FlowJo software (version 10.10).

### Gametocyte culture conditions and standard membrane feeding assay

Sexual conversion in parasites was induced in a trophozoite-stage culture, adding Albumax-containing medium for 36 h and changed every 12 h (65). Gametocytes were then cultured with normal culture medium until maturation. During the entire procedure gametocytes were kept at 37° C in low oxygen conditions. To remove ABS from the culture, gametocytes were cultured in the presence of heparin (20 U/ml) for 4 days. To quantify exflagellation, cultures were mixed 1:1 with 50 μM xanthurenic acid in complete RPMI medium. The solution, kept at room temperature for 15 min, was then transferred to a Neubauer chamber where the exflagellation centres were counted with a 40X objective, bright-field microscopy. To assess mosquito infectivity, day 12 mature gametocytes (65) were added to the bloodmeal of *Anopheles stephensi* mosquitoes (colony maintained at Radboudumc, Nijmegen, The Netherlands), as previously described (66). Mean oocysts per midgut and statistical significance were calculated using a generalized linear mixed effect model with a random experiment effect under a negative binomial distribution. The fitted model was used to obtain estimated means and contrasts that were evaluated using Wald statistics. All datasets, statistical analysis and p-values are reported in Supplementary Information 2.

### Fluorescence microscopy

Mature gametocytes were incubated in 100 nM MitoTracker (MitoTracker® Orange CMTMRos, Thermo Fisher, #M7510) diluted in RPMI medium for exactly 30 min at 37° C and afterwards washed and diluted 1:10 in incomplete medium. The samples were allowed to settle on poly-L-Lysin coated coverslips (Corning, #354085) for 30 min at 37° C before being fixed in 4% paraformaldehyde (Thermo Fisher, #28906) and 0.0075% glutaraldehyde (Panreac, #A0589,0010) in PBS for 20 min (67). The samples were then washed with 1X PBS and permeabilized with 0.1% Triton X-100 in PBS for 10 min. Cells were blocked with 3% BSA for 1 h at RT. The coverslips were incubated with primary antibodies diluted in PBS with 3% BSA (Table S2) for 1 h at RT, washed three times and then incubated with the secondary antibodies for 1 h at RT. All samples were additionally stained with 300 nM DAPI (Thermo Fisher, #62248) for 1 h before being mounted onto microscope slides with Fluoromount G (Thermo Fisher, # 00-4958-02). Images were acquired using an LSM900 confocal microscope with airyscan (ZEISS) using a 63x oil objective with 405, 488, 561, and 633 nm laser excitation and an Electronically Switchable Illumination and Detection Model (ESID) for transmitted light. Images belonging to the same experiment were acquired with equal settings for laser power and detector sensibility and afterwards analyzed with the same parameters to compare expression.

### Mitochondria quantification in fluorescence microscopy

Mature gametocytes prepared as previously explained, were randomly selected based on DIC or DAPI signal to avoid bias. Mitochondria were quantified using the Fiji plug-in MitochondriaAnalyser (36). Briefly, 3D image stacks were pre-processed using the following commands: subtract background (radius = 1.25 μm); sigma filter plus (radius = 2.0 μm); enhance local contrast (slope = 1.40 μm); and gamma correction (value = 0.9 μm). For thresholding, block size was set to 0.85 μm with a C value of 7. Thresholded images were then postprocessed using the functions despeckle and remove outliers with radius = 3 pixels. Statistical significance was determined using one-way ANOVA test with Sidàk correction. Mitochondria were divided in tubular, intermediate and round up based on average branch length (total branch length / number of mitochondria intensities) and statistical significance was determined using a Chi-squared test. To assess membrane potential the same samples were used to quantify MitoTracker signal. The automatic tresholded images were used to determine background-corrected mean fluorescence intensity in the z-stack. All datasets, statistical analysis and p-values are reported in Supplementary Information 2.

### Ultra-Expansion Microscopy (U-ExM)

Mature gametocytes were isolated using MACS LS columns (Miltenyi Biotec). Shortly, LS columns were equilibrated using incomplete RPMI medium and mature gametocyte cultures were flushed in the LS columns attached to a magnet using a 23G needle to regulate the flow. To elute gametocytes, columns were removed from the magnet and washed with incomplete medium. MACS column, magnet and incomplete RPMI medium were previously incubated at 37° C, and kept at this temperature throughout the procedure, to prevent gametocyte activation during purification. The gametocyte pellet was then incubated with 100 nM MitoTracker and fixed as described for fluorescence microscopy. Coverslips were then incubated with 0.5% acrylamide (AA) (Sigma, #A4058) and 0.38% formaldehyde (FA) (Sigma, #F8775) in PBS overnight at 37° C. Gelation was performed as previously described (68). Briefly, the gelling solution was prepared by adding tetramethylethylenediamine (TMED, final concentration 0.5%) (Sigma, #T7024) and ammonium persulfate (APS, final concentration 0.5%) (Sigma, #A3678) to U-ExM monomer solution (MS) (23% w/v sodium acrylate (AK Scientific, #R624), 10% w/v AA, N,N’-methylenbisacrylamide (BIS) (Sigma, #M1533) 0.1% w/v in PBS). The solution was then added to the coverslips and incubated at 37° C for 1 h. Gels were incubated with denaturation buffer (200 mM SDS, 200 mM NaCl, 50 mM Tris in water, pH 9) under strong agitation until complete detachment from the coverslips and transferred into a 1.5 ml Eppendorf tube filled with denaturation buffer and incubated 90 minutes at 95° C. Gels were expanded overnight in ultrapure water at RT. Gels were then incubated with primary antibodies (Table S2) for 3 h at 37° C while shaking, followed by secondary antibodies, NHS-ester (8 μg/ml) (DyLightTM 405 NHS-ester, ThermoScientific, #46400) and SYTOX™ (Thermo Fisher, #S11381) for 2.5 h at 37° C, shaking. For imaging, gels were cut into pieces and positioned on a poly-D lysine (Gibco, #A38904-01) coated coverslips. Images were acquired with identical parameters, using an LSM900 confocal microscope with airyscan (ZEISS) with a 63X oil objective at 405, 488, 561, and 633 nm laser excitation and an Electronically Switchable Illumination and Detection Model (ESID) for transmitted light.

### Electron microscopy sample preparation

Mature gametocytes were harvested using MACS LS columns as described for U-ExM. Purified gametocytes were then fixed with 4% PFA in 0.1 M PHEM and pelleted at 600 x g for 10 min. Pellets were vitrified using the Leica EMPACT high pressure freezer with membrane carrier assembly. Water removal and staining was achieved by freeze substitution in 1% osmium tetroxide, 0.2% uranyl acetate and 1% water in acetone. Stepwise freeze substitution consisted of a 48 h incubation at -90° C, sloped increase to -60° C within 12 h, 10 h incubation at -60° C, sloped increase to -60° C within 12 h, incubation at - 30° C for 8 h and removal of uranyl acetate by washing in 1% osmium tetroxide in acetone at -30° C. The samples were then transferred to ice, osmium tetroxide was replaced with pure acetone, and the samples were further stained with 1% tannic acid in acetone for 1 h and with 1% osmium tetroxide in acetone for 1 h. Then the samples were embedded in Epon (46% HY 964 Araldite, 39.6% Epoxy Embedding Medium, 13% MNA, 1.4% DMP-30) in a stepwise increase of the Epon to acetone ratio. Fully infiltrated samples were hardened at 70° C for 5 days. Thin sections of 65 nm or semi-thin sections of 250 nm were collected on copper grids with formvar/carbon coating. Thin sections were post stained with 2% uranyl acetate for 5 min and Reynolds lead citrate (Reynolds, 1963) for 2 min. Semi-thin sections were labelled with 15 nm gold fiducials for tilt-series alignment.

### Transmission electron tomography

All tomograms were recorded at a JEOL2100 TEM operating at 200 kV. Tomograms were recorded as dual-axis tilt series at 15k magnification, 1.5° tilt increment ranging from -60° to 60° (pixel size 1.52 nm, defocus –1.1 μm). Reconstruction, segmentation and quantifications were performed manually, in IMOD. The statistical significance of the deviations in crista junction diameter was determined using one-way ANOVA test with Sidàk correction. Significance of the difference in crista junction prevalence was tested using a Chi-squared test. All datasets, statistical analysis and p-values are reported in Supplementary Information 2.

### Thin section electron microscopy

Micrographs were collected at a JEOL1400 TEM operating at 120 kV. Recorded images were randomized for blinded quantification of crista length, diameter, area, and coverage. Statistical significance was determined using one-way ANOVA tests with Sidàk correction. All datasets, statistical analysis and p-values are reported in Supplementary Information 2.

### SDS-Page gel and western blot

Saponin pellets of gametocytes and ABS cultures were prepared by resuspending in 7X pellet volume ultrapure water and 2X pellet volume reductive loading buffer (1% β-mercaptoethanol, 0.004% bromophenol blue, 6% glycerol, 2% SDS 50 mM Tris-HCl, pH 6.8). Samples were heated to 70° C for 10 min, quickly vortexed and spun down to precipitate insoluble debris and DNA. The supernatant was loaded on a SurePAGE, 8% Bis-Tris SDS-PAGE (GenScript, #M00677). A standard protein ladder (Precision Plus Protein in Dual Colour Standard, Bio-Rad) was included in every run. Proteins were transferred, with the semi-wet transfer, to a nitrocellulose membrane soaked in transfer buffer (20% methanol with 10% Tris-Glycine (10X)) in a Trans-Blot Turbo machine (Bio-Rad) at 25V for 30 min. The membrane was then blocked in 1% milk powder in PBS with 0.05% Tween 20 (PBST) overnight at 4° C. Primary and secondary antibodies (Table S2) were diluted in 1% milk powder in PBST and incubated for 1 h at RT. The blot was then imaged with Odyssey CLx (ImageQuant).

### Structural comparisons and domain predictions

All AlphaFold models were retrieved from the AlphaFold protein structure database (ScMIC60: AF-P36112-F1-v4, *Pf*MIC60: AF-. Q8IJW5-F1-v4, ScMIC19: AF-P43594-F1-v4, *Pf*MIC19: AF-Q8IIR5-F1-v4)(32, 33). Domains were used as predicted by TED consensus domain, as integrated in the AlphaFold protein structure database (34). Associated pLDDT prediction confidence information is shown in Figure S1. Theoretical length of the coiled coil domain in ScMIC60 and *Pf*MIC60 was measured by atomic distance within the predicted models.

## Supporting information

Response to reviewers

## Acknowledgements

We would like to thank all members of the molecular & cellular parasitology team at the Radboudumc and the microbiology department at the Radboud University for the discussion and support during the project. We would like to thank the General Instrumentation Department at Radboud University, Radboud Technology Center Microscopy and the Radboud Technology Center Flowcytometry for use of their facilities. In addition, we also thank Jordache Ramjith for his advice and support in the statistical analysis and Jolanda Klaassen, Laura Pelser-Posthumus and, Astrid Pouwelsen for breeding of mosquitoes and handling of the infected mosquitoes. We thank Till Voss and Marcus Lee for providing the pBF-gC and pMLB626 CRISPR/Cas9 guide plasmids, respectively. This work was supported by funding provided by the Dutch Research Council (NWO, grant number: OCENW.M.21.087) awarded to Taco W.A. Kooij and Laura van Niftrik. Ultra-expansion microscopy was funded by a Radboud Community of Infectious Disease grant (RCI Grant 2024) awarded to Taco W.A. Kooij.

## Author contributions

LvN and TWAK obtained funding and conceptualized and supervised the project; STL, IB and JLO performed experiments; STL, IB, JLO, FE, TWAK and LvN analysed data; NIP, GJvG and RS executed the mosquito experiments; STL and IB wrote the first draft of the manuscript and all authors reviewed, edited and approved the manuscript.

## Data Availability Statement

All TEM tomography datasets are available in the Electron Microscopy Data Bank (EMDB) under accession codes EMD-54915 (NF54), EMD-55186 (*mic19*^*–*^), EMD-55149 (*mic60*^*–*^) and EMD-55336 (*mic19/mic60*^*–*^).

## References

1. Wurm CA, Jakobs S. Differential protein distributions define two sub-compartments of the mitochondrial inner membrane in yeast. FEBS Lett. 2006;580(24):5628–34.

2. Gilkerson RW, Selker JM, Capaldi RA. The cristal membrane of mitochondria is the principal site of oxidative phosphorylation. FEBS Lett. 2003;546(2-3):355–8.

3. Vogel F, Bornhovd C, Neupert W, Reichert AS. Dynamic subcompartmentalization of the mitochondrial inner membrane. J Cell Biol. 2006;175(2):237–47.

4. Perkins G, Renken C, Martone ME, Young SJ, Ellisman M, Frey T. Electron tomography of neuronal mitochondria: three-dimensional structure and organization of cristae and membrane contacts. J Struct Biol. 1997;119(3):260–72.

5. Muhleip A, Kock Flygaard R, Ovciarikova J, Lacombe A, Fernandes P, Sheiner L, Amunts A. ATP synthase hexamer assemblies shape cristae of Toxoplasma mitochondria. Nat Commun. 2021;12(1):120.

6. Panek T, Elias M, Vancova M, Lukes J, Hashimi H. Returning to the fold for lessons in mitochondrial crista diversity and evolution. Curr Biol. 2020;30(10):R575–R88.

7. Davies KM, Strauss M, Daum B, Kief JH, Osiewacz HD, Rycovska A, et al. Macromolecular organization of ATP synthase and complex I in whole mitochondria. Proc Natl Acad Sci U S A. 2011;108(34):14121–6.

8. Frey TG, Renken CW, Perkins GA. Insight into mitochondrial structure and function from electron tomography. Biochim Biophys Acta. 2002;1555(1-3):196–203.

9. van der Laan M, Horvath SE, Pfanner N. Mitochondrial contact site and cristae organizing system. Curr Opin Cell Biol. 2016;41:33–42.

10. Stephan T, Bruser C, Deckers M, Steyer AM, Balzarotti F, Barbot M, et al. MICOS assembly controls mitochondrial inner membrane remodeling and crista junction redistribution to mediate cristae formation. EMBO J. 2020;39(14):e104105.

11. Hoppins S, Collins SR, Cassidy-Stone A, Hummel E, Devay RM, Lackner LL, et al. A mitochondrial-focused genetic interaction map reveals a scaffold-like complex required for inner membrane organization in mitochondria. J Cell Biol. 2011;195(2):323–40.

12. Harner M, Korner C, Walther D, Mokranjac D, Kaesmacher J, Welsch U, et al. The mitochondrial contact site complex, a determinant of mitochondrial architecture. EMBO J. 2011;30(21):4356–70.

13. von der Malsburg K, Muller JM, Bohnert M, Oeljeklaus S, Kwiatkowska P, Becker T, et al. Dual role of mitofilin in mitochondrial membrane organization and protein biogenesis. Dev Cell. 2011;21(4):694–707.

14. Huynen MA, Muhlmeister M, Gotthardt K, Guerrero-Castillo S, Brandt U. Evolution and structural organization of the mitochondrial contact site (MICOS) complex and the mitochondrial intermembrane space bridging (MIB) complex. Biochim Biophys Acta. 2016;1863(1):91–101.

15. Zerbes RM, Bohnert M, Stroud DA, von der Malsburg K, Kram A, Oeljeklaus S, et al. Role of MINOS in mitochondrial membrane architecture: cristae morphology and outer membrane interactions differentially depend on mitofilin domains. J Mol Biol. 2012;422(2):183–91.

16. Friedman JR, Mourier A, Yamada J, McCaffery JM, Nunnari J. MICOS coordinates with respiratory complexes and lipids to establish mitochondrial inner membrane architecture. Elife. 2015;4.

17. Tarasenko D, Barbot M, Jans DC, Kroppen B, Sadowski B, Heim G, et al. The MICOS component Mic60 displays a conserved membrane-bending activity that is necessary for normal cristae morphology. J Cell Biol. 2017;216(4):889–99.

18. Barbot M, Jans DC, Schulz C, Denkert N, Kroppen B, Hoppert M, et al. Mic10 oligomerizes to bend mitochondrial inner membranes at cristae junctions. Cell Metab. 2015;21(5):756–63.

19. Bock-Bierbaum T, Funck K, Wollweber F, Lisicki E, von der Malsburg K, von der Malsburg A, et al. Structural insights into crista junction formation by the Mic60-Mic19 complex. Sci Adv. 2022;8(35):eabo4946.

20. Rabl R, Soubannier V, Scholz R, Vogel F, Mendl N, Vasiljev-Neumeyer A, et al. Formation of cristae and crista junctions in mitochondria depends on antagonism between Fcj1 and Su e/g. J Cell Biol. 2009;185(6):1047–63.

21. Korner C, Barrera M, Dukanovic J, Eydt K, Harner M, Rabl R, et al. The C-terminal domain of Fcj1 is required for formation of crista junctions and interacts with the TOB/SAM complex in mitochondria. Mol Biol Cell. 2012;23(11):2143–55.

22. Tassan-Lugrezin S. DSACA, van Niftrik L, Kooij T. W. A., Bregy I. Unconventional model organisms bend our view on mitochondrial cristae. Journal of Cell science, in press. 2026.

23. Evers F, Cabrera-Orefice A, Elurbe DM, Kea-Te Lindert M, Boltryk SD, Voss TS, et al. Composition and stage dynamics of mitochondrial complexes in Plasmodium falciparum. Nat Commun. 2021;12(1):3820.

24. Nina PB, Morrisey JM, Ganesan SM, Ke H, Pershing AM, Mather MW, Vaidya AB. ATP synthase complex of Plasmodium falciparum: dimeric assembly in mitochondrial membranes and resistance to genetic disruption. J Biol Chem. 2011;286(48):41312–22.

25. Evers F, Roverts R, Boshoven C, Kea-Te Lindert M, Verhoef JMJ, Sommerdijk N, et al. Comparative 3D ultrastructure of Plasmodium falciparum gametocytes. Nat Commun. 2025;16(1):69.

26. Ke H, Lewis IA, Morrisey JM, McLean KJ, Ganesan SM, Painter HJ, et al. Genetic investigation of tricarboxylic acid metabolism during the Plasmodium falciparum life cycle. Cell Rep. 2015;11(1):164–74.

27. MacRae JI, Dixon MW, Dearnley MK, Chua HH, Chambers JM, Kenny S, et al. Mitochondrial metabolism of sexual and asexual blood stages of the malaria parasite Plasmodium falciparum. BMC Biol. 2013;11:67.

28. Srivastava A, Philip N, Hughes KR, Georgiou K, MacRae JI, Barrett MP, et al. Stage-specific changes in Plasmodium metabolism required for differentiation and adaptation to different host and vector environments. PLoS Pathog. 2016;12(12):e1006094.

29. Sheokand PK, Pradhan S, Maclean AE, Muhleip A, Sheiner L. Plasmodium falciparum Mitochondrial Complex III, the Target of Atovaquone, Is Essential for Progression to the Transmissible Sexual Stages. Int J Mol Sci. 2024;25(17).

30. Chen F, Mackey AJ, Stoeckert CJ, Jr., Roos DS. OrthoMCL-DB: querying a comprehensive multi-species collection of ortholog groups. Nucleic Acids Res. 2006;34(Database issue):D363–8.

31. Verhoef JMJ, Boshoven C, Evers F, Akkerman LJ, Gijsbrechts BCA, van de Vegte-Bolmer M, et al. Detailing organelle division and segregation in Plasmodium falciparum. J Cell Biol. 2024;223(12).

32. Jumper J, Evans R, Pritzel A, Green T, Figurnov M, Ronneberger O, et al. Highly accurate protein structure prediction with AlphaFold. Nature. 2021;596(7873):583–9.

33. Varadi M, Anyango S, Deshpande M, Nair S, Natassia C, Yordanova G, et al. AlphaFold Protein Structure Database: massively expanding the structural coverage of protein-sequence space with high-accuracy models. Nucleic Acids Res. 2022;50(D1):D439–D44.

34. Lau AM, Bordin N, Kandathil SM, Sillitoe I, Waman VP, Wells J, et al. Exploring structural diversity across the protein universe with The Encyclopedia of Domains. Science. 2024;386(6721):eadq4946.

35. Rawlings DJ, Fujioka H, Fried M, Keister DB, Aikawa M, Kaslow DC. Alpha-tubulin II is a male-specific protein in Plasmodium falciparum. Mol Biochem Parasitol. 1992;56(2):239–50.

36. Chaudhry A, Shi R, Luciani DS. A pipeline for multidimensional confocal analysis of mitochondrial morphology, function, and dynamics in pancreatic beta-cells. Am J Physiol Endocrinol Metab. 2020;318(2):E87–E101.

37. Schindelin J, Arganda-Carreras I, Frise E, Kaynig V, Longair M, Pietzsch T, et al. Fiji: an open-source platform for biological-image analysis. Nat Methods. 2012;9(7):676–82.

38. Anand R, Reichert AS, Kondadi AK. Emerging roles of the MICOS complex in cristae dynamics and biogenesis. Biology (Basel). 2021;10(7).

39. Decelle J, Kayal E, Bigeard E, Gallet B, Bougoure J, Clode P, et al. Intracellular development and impact of a marine eukaryotic parasite on its zombified microalgal host. The ISME Journal. 2022;16(10):2348–59.

40. Sturm A, Mollard V, Cozijnsen A, Goodman CD, McFadden GI. Mitochondrial ATP synthase is dispensable in blood-stage Plasmodium berghei rodent malaria but essential in the mosquito phase. Proc Natl Acad Sci U S A. 2015;112(33):10216–23.

41. Goodman CD, Buchanan HD, McFadden GI. Is the mitochondrion a good malaria drug target? Trends in Parasitology. 2017;33(3):185–93.

42. Sparkes PC, Famodimu MT, Alves E, Springer E, Przyborski J, Delves MJ. Mitochondrial ATP synthesis is essential for efficient gametogenesis in Plasmodium falciparum. Commun Biol. 2024;7(1):1525.

43. Otto TD, Wilinski D, Assefa S, Keane TM, Sarry LR, Bohme U, et al. New insights into the blood-stage transcriptome of Plasmodium falciparum using RNA-Seq. Mol Microbiol. 2010;76(1):12–24.

44. Lopez-Barragan MJ, Lemieux J, Quinones M, Williamson KC, Molina-Cruz A, Cui K, et al. Directional gene expression and antisense transcripts in sexual and asexual stages of Plasmodium falciparum. BMC Genomics. 2011;12:587.

45. Weber TA, Koob S, Heide H, Wittig I, Head B, van der Bliek A, et al. APOOL is a cardiolipin-binding constituent of the Mitofilin/MINOS protein complex determining cristae morphology in mammalian mitochondria. PLoS One. 2013;8(5):e63683.

46. Kaurov I, Vancova M, Schimanski B, Cadena LR, Heller J, Bily T, et al. The diverged Trypanosome MICOS complex as a hub for mitochondrial cristae shaping and protein import. Curr Biol. 2018;28(21):3393–407 e5.

47. Kondadi AK, Anand R, Hansch S, Urbach J, Zobel T, Wolf DM, et al. Cristae undergo continuous cycles of membrane remodelling in a MICOS-dependent manner. EMBO Rep. 2020;21(3):e49776.

48. Sheikh S, Turpin Knotková B, Benz C, Eliáš M, Bílý T, Bondar A, et al. The core MICOS complex subunit Mic60 has been substituted by two cryptic mitofilin-containing proteins in Euglenozoa. Mol Biol Evol. 2025;2025 Nov 10:msaf289.

49. Rampelt H, Wollweber F, Licheva M, de Boer R, Perschil I, Steidle L, et al. Dual role of Mic10 in mitochondrial cristae organization and ATP synthase-linked metabolic adaptation and respiratory growth. Cell Rep. 2022;38(4):110290.

50. Cadena LR, Gahura O, Panicucci B, Zikova A, Hashimi H. Mitochondrial contact site and cristae organization system and F(1)F(O)-ATP synthase crosstalk is a fundamental property of mitochondrial cristae. mSphere. 2021;6(3):e0032721.

51. Kubihal S, Goyal A, Gupta Y, Khadgawat R. Glucose measurement in body fluids: A ready reckoner for clinicians. Diabetes Metab Syndr. 2021;15(1):45–53.

52. Benning FMC, Bell TA, Nguyen TH, Syau D, Connell LB, Liao YT, et al. Ancestral sequence reconstruction of the Mic60 Mitofilin domain reveals residues supporting respiration in yeast. Protein Sci. 2025;34(7):e70207.

53. Okamoto N, Spurck TP, Goodman CD, McFadden GI. Apicoplast and mitochondrion in gametocytogenesis of Plasmodium falciparum. Eukaryot Cell. 2009;8(1):128–32.

54. Frey TG, Mannella CA. The internal structure of mitochondria. Trends in Biochemical Sciences. 2000;25(7):319–24.

55. Nicastro D, Frangakis AS, Typke D, Baumeister W. Cryo-electron tomography of neurospora mitochondria. J Struct Biol. 2000;129(1):48–56.

56. Perkins GA, Ellisman MH, Fox DA. Three-dimensional analysis of mouse rod and cone mitochondrial cristae architecture: bioenergetic and functional implications. Mol Vis. 2003;9:60–73.

57. Verhoef JMJ, Bekkering ET, Boshoven C, Hannon M, Evers F, Proellochs NI, et al. The role of stomatin-like protein (STOML) in Plasmodium falciparum. bioRxiv. 2024:2024.07.18.604071.

58. Kooij TW, Rauch MM, Matuschewski K. Expansion of experimental genetics approaches for Plasmodium berghei with versatile transfection vectors. Mol Biochem Parasitol. 2012;185(1):19–26.

59. Filarsky M, Fraschka SA, Niederwieser I, Brancucci NMB, Carrington E, Carrio E, et al. GDV1 induces sexual commitment of malaria parasites by antagonizing HP1-dependent gene silencing. Science. 2018;359(6381):1259–63.

60. Lim MY, LaMonte G, Lee MCS, Reimer C, Tan BH, Corey V, et al. UDP-galactose and acetyl-CoA transporters as Plasmodium multidrug resistance genes. Nat Microbiol. 2016;1:16166.

61. Trager W, Jensen JB. Human malaria parasites in continuous culture. Science. 1976;193(4254):673–5.

62. Lambros C, Vanderberg JP. Synchronization of Plasmodium falciparum erythrocytic stages in culture. The Journal of Parasitology. 1979;65(3):418–20.

63. Wu Y, Sifri CD, Lei HH, Su XZ, Wellems TE. Transfection of Plasmodium falciparum within human red blood cells. Proc Natl Acad Sci U S A. 1995;92(4):973–7.

64. Kenthirapalan S, Waters AP, Matuschewski K, Kooij TW. Flow cytometry-assisted rapid isolation of recombinant Plasmodium berghei parasites exemplified by functional analysis of aquaglyceroporin. Int J Parasitol. 2012;42(13-14):1185–92.

65. Graumans W, van der Starre A, Stoter R, van Gemert GJ, Andolina C, Ramjith J, et al. AlbuMAX supplemented media induces the formation of transmission-competent P. falciparum gametocytes. Mol Biochem Parasitol. 2024;259:111634.

66. Stone WJ, Eldering M, van Gemert GJ, Lanke KH, Grignard L, van de Vegte-Bolmer MG, et al. The relevance and applicability of oocyst prevalence as a read-out for mosquito feeding assays. Sci Rep. 2013;3:3418.

67. Tonkin CJ, van Dooren GG, Spurck TP, Struck NS, Good RT, Handman E, et al. Localization of organellar proteins in Plasmodium falciparum using a novel set of transfection vectors and a new immunofluorescence fixation method. Mol Biochem Parasitol. 2004;137(1):13–21.

68. Gambarotto D, Hamel V, Guichard P. Ultrastructure expansion microscopy (U-ExM). Methods Cell Biol. 2021;161:57–81.

