## Supplementary material for "The role of MICOS in organizing mitochondrial cristae in malaria parasites": Response to reviewers

Reviewer #1 (Evidence, reproducibility and clarity (Required)): 

Summary: This manuscript reports the identification of putative orthologues of mitochondrial contact site and cristae organizing system (MICOS) proteins in Plasmodium falciparum - an organism that unusually shows an acristate mitochondrion during the asexual part of its life cycle and then this develops cristae as it enters the sexual stage of its life cycle and beyond into the mosquito. The authors identify PfMIC60 and PfMIC19 as putative members and study these in detail. The authors at HA tags to both proteins and look for timing of expression during the parasite life cycle and attempt (unsuccessfully) to localise them within the parasite. They also genetically deleted both gene singly and in parallel and phenotyped the effect on parasite development. They show that both proteins are expressed in gametocytes and not asexuals, suggesting they are present at the same time as cristae development. They also show that the proteins are dispensible for the entire parasite life cycle investigated (asexuals through to sporozoites), however there is some reduction in mosquito transmission. Using EM techniques they show that the morphology of gametocyte mitochondria is abnormal in the knock out lines, although there is great variation.

Major comments: The manuscript is interesting and is an intriguing use of a well studied organism of medical importance to answer fundamental biological questions. My main comments are that there should be greater detail in areas around methodology and statistical tests used. Also, the mosquito transmission assays (which are notoriously difficult to perform) show substantial variation between replicates and the statistical tests and data presentation are not clear enough to conclude the reduction in transmission that is claimed. Perhaps this could be improved with clearer text?

We would like to thank the reviewer for taking the time to review our manuscript. We are happy to hear the reviewer thinks the manuscript is interesting and thank the reviewer for their constructive feedback.

To clarify the statistical analyses used, we included a new supplementary dataset with all statistical analyses and p-values indicated per graph. Furthermore, figure legends now include the information on the exact statistical test used in each case.

Regarding mosquito experiments, while we indeed reported a reduction in transmission and oocysts numbers we are aware that this effect might be due to the high variability in mosquito feeding assays. To highlight this point, we deleted the sentence “with the transmission reduction of [numbers]….” and we included the sentence “The high variability encountered in the standard membrane feeding assays, though, partially obstructs a clear conclusion on the biological relevance of the observed reduction in oocyst numbers“

More specific comments to address: 
Line 101/Fig1E (and figure legend) - What is this heatmap showing. It would be helpful to have a sentence or two linking it to a specific methodology. I could not find details in the M+M section and "specialized, high molecular mass gels" does not adequately explain what experiments were performed. The reference to Supplementary Information 1 also did not provide information.

We added the information “high molecular mass gels with lower acrylamide percentage” to clarify methodology in the text. Furthermore, we extended the figure legend to include all relevant information. Further experimental details can be found in the study cited in this context, where the dataset originates from (Evers et al., 2021).

Line 115 and Supplementary Figure 2C + D - The main text says that the transgenic parasites contained a mitochondrially localized mScarlet for visualization and localization, but in the supplementary figure 2 it shows mitotracker labelling rather than mScarlet. This is very confusing. The figure legend also mentions both mScarlet and MitoTracker. I assume that mScarlet was used to view in regular IFAs (Fig S2C) and the MitoTracker was used for the expansion microscopy (Fig S2D)? Please clarify.

We thank the reviewer for pointing this out – this was indeed incorrectly annotated. We used the endogenous mito-mScarlet signal in IFA and mitoTracker in U-ExM. The figure annotation has now been corrected.

Figure 2C - what is the statistical test being used (the methods say "Mean oocysts per midgut and statistical significance were calculated using a generalized linear mixed effect model with a random experiment effect under a negative binomial distribution." but what test is this?)? The statistic test is now included in the material and method section with the sentence “The fitted model was used to obtain estimated means and contrasts and were evaluated using Wald Statistics”. The test is now also mentioned in the figure legend.

Also the choice of a log10 scale for oocyst intensity is an unusual choice - how are the mosquitoes with 0 oocysts being represented on this graph? It looks like they are being plotted at 10^-1 (which would be 0.1 oocysts in a mosquito which would be impossible).

As the data spans three orders of magnitude with low values being biologically meaningful, we decided that a log scale would best facilitate readability of the graph. As the 0 values are also important to show, we went with a standard approach to handle 0s in log transformed data and substituted the 0s with a small value (0.001). We apologize for not mentioning this transformation in the manuscript. To make this transformation transparent, we added a break at the lower end of the log‑scaled y‑axis and relabelled the lowest tick as ‘0’. This ensures that mosquitoes with zero oocysts are shown along the x‑axis without being assigned an artificial value on the log scale. We would furthermore like to highlight that for statistics we used the true value 0 and not 0.001.

Figure 2D - it is great that the data from all feeding replicates has been shared, however it is difficult to conclude any meaningful impact in transmission with the knock-out lines when there is so much variation and so few mosquitoes dissected for some datapoints (10 mosquitoes are very small sample sizes). For example, Exp1 shows a clear decrease in mic19- transmission, but then Exp2 does not really show as great effect. Similarly, why does the double knock out have better transmission than the single knockouts? Sure there would be a greater effect?

We agree with the reviewer and with the new sentence added, as per major point, we hope we clarified the concept. Note that original Figure 2D has been moved to the supplementary information, as per minor comment of another reviewer.

Figure 3 legend - Please add which statistical test was used and the number of replicates.

Done

Figure 4 legend - Please add which statistical test was used and the number of replicates.

Done. Regarding replicates, note that while we measured over 100 cristae from over 30 mitochondria, these all stem from the same parasite culture.

Figure 5C - the 3D reconstructions are very nice, but what does the red and yellow coloring show?

Indeed, the information was missing. We added it to the figure legend.

Line 352 - "Still, it is striking that, despite the pronounced morphological phenotype, and the possibly high mitochondrial stress levels, the parasites appeared mostly unaffected in life cycle propagation, raising questions about the functional relevance of mitochondria at these stages." 
How do the authors reconcile this statement with the proven fact that mitochondria-targeted antimalarials (such as atovaquone) are very potent inhibitors of parasite mosquito transmission? 
Our original sentence was reductive. What we wanted to state was related to the functional relevance of crista architecture and overall mitochondrial morphology rather than the general functional relevance of the mitochondria. We changed the sentence accordingly.

Furthermore, even though we do not discuss this in the article, we are aware of mitochondria targeting drugs that are known to block mosquito transmission. We want to point out that it is difficult to discern the disruption of ETC and therefore an impact on energy conversion with the impact on the essential pathway of pyrimidine synthesis, highly relevant in microgamete formation. Still, a recent paper from Sparkes et al. 2024 showed the essentiality of mitochondrial ATP synthesis during gametogenesis so it is very likely that the mitochondrial energy conversion is highly relevant for transmission to the mosquito.

Reviewer #1 (Significance (Required)): 

This manuscript is a novel approach to studying mitochondrial biology and does open a lot of unanswered questions for further research directions. Currently there are limitations in the use of statistical tests and detail of methodology, but these could be easily be addressed with a bit more analysis/better explanation in the text. 
This manuscript could be of interest to readers with a general interest in mitochondrial cell biology and those within the specific field of Plasmodium research. 
My expertise is in Plasmodium cell biology. 
We thank the reviewer for the praise.

Reviewer #2 (Evidence, reproducibility and clarity (Required)): 

Major comments: 
1) In my opinion, the authors tend to sensationalize or overinterpret their results. The title of the manuscript is very misleading. While MICOS is certainly important for crista formation, it is not the only factor, as ATP synthase dimer rows make a highly significant contribution to crista morphology. Thus, one can argue with equal validity that ATP synthase should be considered the 'architect', as it's the conformation of the dimers and rows modulate positive curvature. Secondly, while cristae are still formed upon mic60/mic19 gene knockout (KO), they are severely deformed, and likely dysfunctional (see below). Thus, I do not agree with the title that MICOS is dispensable for crista formation, because the authors results show that it clearly is essential. So, the title should be changed.

We thank the reviewer for taking the time to review our manuscript.

Based on the reviewers’ interpretation we conclude the title does not come across as intended. We have changed the title to: “The role of MICOS in organizing mitochondrial cristae in malaria parasites”

The Discussion section starting from line 373 also suffers from overinterpretation as well as being repetitive and hard to understand. The authors infer that MICOS stability is compromised less in the single KOs (sKO) in compared to the mic60/mic19 double KO (dKO). MICOS stability was never directly addressed here and the composition of the MICOS complex is unaddressed, so it does not make sense to speculate by such tenuous connections. The data suggest to me that mic60 and mic19 are equally important for crista formation and crista junction (CJ) stabilization, and the dKO has a more severe phenotype than either KO, further demonstrating neither is epistatic.

We do agree with the reviewer’s notion that we did not address complex stability, and our wording did not make this sufficiently clear. We shortened and rephrased the paragraph in question.

The following paragraphs (line 387 to 422) continues with such unnecessary overinterpretation to the point that it is confusing and contradictory. Line 387 mentions an 'almost complete loss of CJs' and then line 411 mentions an increase in CJ diameter, both upon Mic60 ablation. I do not think this discussion brings any added value to the manuscript and should be shortened. Yes, maybe there are other putative MICOS subunits that may linger in the KOS that are further destabilized in the dKO, or maybe Mic60 remains in the mic19 KO (and vice versa) to somehow salvage more CJs, which is not possible in the dKO. It is impossible to say with confidence how ATP synthase behaves in the KOs with the current data. 
We shortened this paragraph.

2) While the authors went through impressive lengths to detect any effect on lifecycle progression, none was found except for a reduction in oocyte count. However, the authors did not address any direct effect on mitochondria, such as OXPHOS complex assembly, respiration, membrane potential. This seems like a missed opportunity, given the team's previous and very nice work mapping these complexes by complexome profiling. However, I think there are some experiments the authors can still do to address any mitochondrial defects using what they have and not resorting to complexome profiling (although this would be definitive if it is feasible):


i) Quantification of MitoTracker Red staining in WT and KOs. The authors used this dye to visualize mitochondria to assay their gross morphology, but unfortunately not to assay membrane potential in the mutants. The authors can compare relative intensities of the different mitochondria types they categorized in Fig. 3A in 20-30 cells to determine if membrane potential is affected when the cristae are deformed in the mutants. One would predict they are affected.

Interesting suggestion. As our staining and imaging conditions are suitable for such analysis (as demonstrated by Sarazin et al., 2025, <https://www.biorxiv.org/content/10.1101/2025.11.27.690934v1>), we performed the measurements on the same dataset which we collected for Figure 3. We did, however, not detect any difference in mitotracker intensity between the different lines. The result of this analysis is included in the new version of Supplementary figure S6.

ii) Sporozoites are shown in Fig S5. The authors can use the same set up to track their motion, with the hypothesis that they will be slower in the mutants compared to WT due to less ATP. This assumes that sporozoite mitochondria are active as in gametocytes.

While theoretically plausible and informative, we currently do not know the relevance of mitochondrial energy conversion for general sporozoite biology or specifically features of sporozoite movement. Given the required resources and time to set this experiment up and the uncertainty whether it is a relevant proxy for mitochondrial functioning, we argue it is out of scope for this manuscript.

iii) Shotgun proteomics to compare protein levels in mutants compared to WT, with the hypothesis that OXPHOS complex subunits will be destabilized in the mutants with deformed cristae. This could be indirect evidence that OXPHOS assembly is affected, resulting in destabilized subunits that fail to incorporate into their respective complexes.

While this experiment could potentially further our understanding of the interaction between MICOS and levels of OXPHOS complex subunits we argue that the indirect nature of the evidence does not justify the required investments.

To expedite resubmission, the authors can restrict the cell lines to WT and the dKO, as the latter has a stronger phenotype that the individual KOs and conclusions from this cell line are valid for overall conclusions about Plasmodium MICOS.

I will also conclude that complexome/shotgun proteomics may be a useful tool also for identifying other putative MICOS subunits by determining if proteins sharing the same complexome profile as PfMic60 and Mic19 are affected. This would address the overinterpretation problem of point 1. 

3) I am aware of the authors previous work in which they were not able to detect cristae in ABS, and thus have concluded that these are truly acristate. This can very well be true, or there can be immature cristae forms that evaded detection at the resolution they used in their volumetric EM acquisitions. The mitochondria and gametocyte cristae are pretty small anyway, so it not unreasonable to assume that putative rudimentary cristae in ABS may be even smaller still. Minute levels of sampled complex III and IV plus complex V dimers in ABS that were detected previously by the authors by complexome profiling would argue for the presence of miniscule and/or very few cristae.

I think that authors should hedge their claim that ABS is acrisate by briefly stating that there still is a possibility that miniscule cristae may have been overlooked previously.
We acknowledge that we cannot demonstrate the absolute absence of any membrane irregularities along the inner mitochondrial membrane. At the same time, if such structures were present, they would be extremely small and unlikely to contain the full set of proteins characteristic of mature cristae. For this reason, we consider it appropriate to classify ABS mitochondria as acristate. To reflect the reviewer’s point while maintaining clarity for readers, we have slightly adjusted our wording in the manuscript, changing ‘fully acristate’ to ‘acristate’.

This brings me to the claim that Mic19 and Mic60 proteins are not expressed in ABS. This is based on the lack of signal from the epitope tag; a weak signal is detected in gametocytes. Thus, one can counter that Mic19 and Mic60 are also expressed, but below the expression limits of the assay, as the protein exhibits low expression levels when mitochondrial activity is upregulated.

We agree with the reviewer that the absence of a detectable epitope‑tag signal does not definitively exclude low‑level expression, and we have therefore replaced the term ‘absent’ with ‘undetectable’ throughout the manuscript. In context with previous findings of low-level transcripts of the proteins in a study by Lopez-Berragan et al. and Otto et al., we also added the sentence “The apparent absence could indicate that transcripts are not translated in ABS or that the proteins’ expression was below detection limits of western blot analysis.” to the discussion. _At the same time, we would like to clarify that transcript levels for both genes fall within the <25th percentile, suggesting that these low values likely represent background signal rather than biologically meaningful expression. This interpretation is further supported by proteomic datasets in PlasmoDB, which report PfMIC19 and PfMIC60 expression in gametocyte and mosquito stages, but not in asexual blood stages.”

To address this point, the authors should determine of mature mic60 and mic19 mRNAs are detected in ABS in comparison to the dKO, which will lack either transcript. RT-qPCR using polyT primers can be employed to detect these transcripts. If the level of these mRNAs are equivalent to dKO in WT ABS, the authors can make a pretty strong case for the absence of cristae in ABS.

We appreciate the reviewer’s suggestion. As noted in the Discussion, existing transcriptomic datasets already show detectable MIC19 and MIC60 mRNAs in ABS. For this reason, we expect RT-qPCR to reveal low (but not absent) levels of both transcripts, unlike the true loss expected to be observed in the dKO. Because such residual signals have been reported previously and their biological relevance remains uncertain, we do not believe transcript levels alone can serve as a definitive indicator of cristae absence in ABS.

They should highlight the twin CX9C motifs that are a hallmark of Mic19 and other proteins that undergo oxidative folding via the MIA pathway. Interestingly, the Mia40 oxidoreductase that is central to MIA in yeast and animals, is absent in apicomplexans (DOI: 10.1080/19420889.2015.1094593).

Searching for the CX9C motifs is a valuable suggestion. In response to the reviewer´s suggestion we analysed the conservation of the motif in PfMIC19 and included this in a new figure panel (Figure 1 F).

Did the authors try to align Plasmodium Mic19 orthologs with conventional Mic19s? This may reveal some conserved residues within and outside of the CHCH domain.

In response to this comment we made Figure 1 F, where we show conserved residues within the CHCH domains of a broad range of MIC19 annotated sequences across the opisthokonts, and show that the Cx9C motifs are conserved also in PfMIC19. Outside the CHCH domain, we did not find any meaningful conservation, as PfMIC19 heavily diverges from opisthokont MIC19.

5) Statistcal significance. Sometimes my eyes see population differences that are considered insignificant by the statistical methods employed by the authors, eg Fig. 4E, mutants compared to WT, especially the dKO. Have the authors considered using other methods such as student t-test for pairwise comparisons?

The graphs in figures 3, 4 and 5 got a makeover, such that they now are in linear scale and violin plots (also following a suggestion from further down in the reviewer’s comments). We believe that this improves interpretability. ANOVA was kept as statistical testing to assure the correction for multiple comparisons that cannot be performed with standard t-test. A full overview of statistics and exact p-values can also be found in the newly added supplementary information 2.

Minor comments:
Line 33. Anaerobes (eg Giardia) have mitochondria that do produce ATP, unlike aerobic mitochondria
We acknowledge that producing ATP via OXPHOS is not a characteristic of all mitochondria-like organelles (e.g. mitosomes), which is why these are typically classified separately from canonical mitochondria. When not considering mitochondria-like organelles, energy conversion is the function that the mitochondrion is most well-known for and the one associated with cristae.

Line 56: Unclear what authors mean by "canonical model of mitochondria"
To clarify we changed this to “yeast or human” model of mitochondria.

Lines 75-76: This applies to Mic10 only
We removed the “high degree of conservation in other cristate eukaryotes” statement.

Line 80: Cite DOI: 10.1016/j.cub.2020.02.053

Done


Fig 2D: I find this table difficult to read. If authors keep table format, at least get rid of 'mean' column' as this data is better depicted in 2C. I suggest depicted this data either like in 3B depicting portion of infected vs unaffected flies in all experiments, then move modified Table to supplement. Important to point out experiment 5 appears to be an outlier with reduced infectivity across all cell lines, including WT.

To clarify: the mean reported in the table indicates the mean per replicate while the mean reported in figure 2C is the overall mean for a given genotype that corrects for variability within experiments. We agree that moving the table to the supplementary data is a good idea. We decided to not include a graph for infected and non-infected mosquitoes as this information would be partially misleading, highlighting a phenotype we argue to be influenced by the strong variability.

Fig. 3C-G: I feel like these data repeatedly lead to same conclusions. These are all different ways of showing what is depicted in Fig 2B: mitochondria gross morphology is affected upon ablation of MICOS. I suggest that these graphs be moved to supplement and replaced by the beautiful images.

Thank you for the nice comment on our images. We have now moved part of the graphs to supplementary figure 6 and only kept the Relative Frequency, Sphericity and total mitochondria volume per cell in the main figure.

Line 180: Be more specific with which tubulin isoform is used as a male marker and state why this marker was used in supplemental Fig S6.

We have now specified the exact tubulin isoform used as the male gametocyte marker, both in the main text and in Supplementary Fig. S6. This is a commercial antibody previously known to work as an effective male marker, which is why we selected it for this experiment. This is now clearly stated in the manuscript.

Line 196 and Fig 3C: the word 'intensities' in this context is very ambiguous. Please choose a different term (puncta, elements, parts?). This is related to major point 2i above.

To clarify the biological effect that we can conclude form the measurement, we added an explanation about it in the respective section of the results, and we decided to replace the raw results of the plug-in readout with the deduced relative dispersion.

Line 222: Report male/female crista measurements
We added Supplementary information 2, which contains exact statistical test and outcomes on all presented quantifications as well as a per-sex statistical analysis of the data from figure 4. Correspondingly, we extended supplementary information 2 by a per-sex colour code for the thin section TEM data.

Fig. 4B-E: depict data as violin plots or scatter plots like Fig. 2C to get a better grasp of how the crista coverage is distributed. It seems like the data spread is wider in the double KO. This would also solve the problem with the standard deviation extending beyond 0%.
We changed this accordingly.

Lines 331-333: Please clarify that this applies for some, but not all MICOS subunits. Please also see major point 1 above. Also, the authors should point out that despite their structural divergence, trypanosomal cryptic mitofilins Mic34 and Mic40 are essential for parasite growth, in contrast to their findings with PfMic60 (DOI: <https://doi.org/10.1101/2025.01.31.635831>).

This has been changed accordingly.


Line 320: incorrect citation. Related to point 1above.

Correct citation is now included in the text.

Lines 333-335. This is related to the above. Again, some subunits appear to affect cell growth under lab conditions, and some do not. This and the previous sentence should be rewritten to reflect this.

This has been changed accordingly.

Line 343-345: The sentence and citation 45 are strange. Regarding the former, it is about CHCHD10, whose status as a bona fide MICOS subunit is very tenuous, so I would omit this. About the phenomenon observed, I think it makes more sense to write that Mic60 ablation results in partially fragmented mitochondria in yeast (Rabl et al., 2009 J Cell Biol. 185: 1047-63). A fragmented mitochondria is often a physiological response to stress. I would just rewrite as not to imply that mitochondrial fission (or fusion) is impaired in these KOs, or at least this could be one of several possibilities.

The sentence has been substituted following the indication of the reviewer. Though we still include the data of the human cells as this has also been shown in Stephens et al. 2020.

Line 373: 'This indicates' is too strong. I would say 'may suggest' as you have no proof that any of the KOs disrupts MICOS. This hypothesis can be tested by other means, but not by penetrance of a phenotype. 
Done

Line 376-377; 'deplete functionality' does not make sense, especially in the context of talking about MICOS subunit stability. In my opinion, this paragraph overinterprets the KO effects on MICOS stability. None of the experiments address this phenomenon, and thus the authors should not try to interpret their results in this context. See major point 1. 
Other suggestions for added value
We removed the sentence. Also, the entire paragraph has been shortened, restructured and wording was changed to address major point 1.

1) Does Plasmodium Sam50 co-fractionate with Mic60 and Mic19 in BN PAGE (Fig. 1E)

While we did identify SAMM50 in our BN PAGE, the protein does not co-migrate with the MICOS components but instead comigrates with other components of a putative sorting and assembly machinery (SAM) complex. As SAMM50, the SAM complex and the overarching putative mitochondrial membrane space bridging (MIB) complex are not mentioned in the manuscript, we decided to not include the information in the figure.

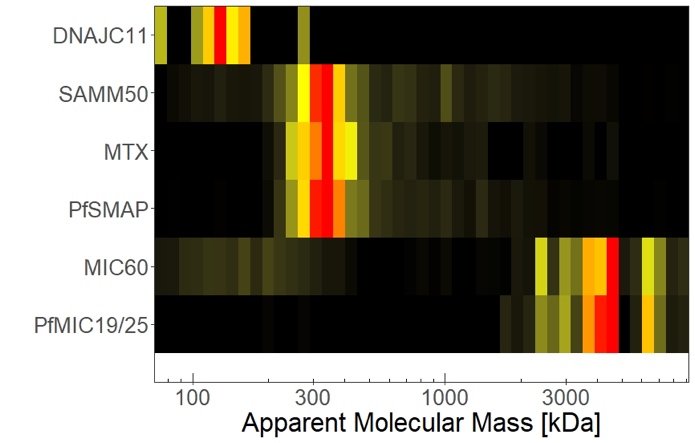

Reviewer #2 (Significance (Required)): 

The manuscript by Tassan-Lugrezin is predicated on the idea that Plasmodium represents the only system in which de novo crista formation can be studied. They leverage this system to ask the question whether MICOS is essential for this process. They conclude based on their data that the answer is no, which the authors consider unprecedented. But even if their claim is true that ABS is acristate, this supposed advantage does not really bring any meaningful insight into how MICOS works in Plasmodium.

First the positives of this manuscript. As has been the case with this research team, the manuscript is very sophisticated in the experimental approaches that are made. The highlights are the beautiful and often conclusive microscopy performed by the authors. Only the localization of Mic60 and Mic19 was inconclusive due to their very low expression unfortunately. 

The examination of the MICOS mutants during in vitro life cycle of Plasmodium falciparum is extremely impressive and yields convincing results. Mitochondrial deformation is tolerated by life cycle stage differentiation, with a modest but significant reduction of oocyte production, being observed. 

However, despite the herculean efforts of the authors, the manuscript as it currently stands represents only a minor advance in our understanding of the evolution of MICOS, which from the title and focus of the manuscript, is the main goal of the authors. 
In its current form, the manuscript reports some potentially important findings: 

1) Mic60 is verified to play a role in crista formation, as is predicted by its orthology to other characterized Mic60 orthologs.

2) The discovery of a novel Mic19 analog (since the authors maintain there is no significant sequence homology), which exhibits a similar (or the same?) complexome profile with Mic60. This protein was upregulated in gametocytes like Mic60 and phenocopies Mic60 KO.

3) Both of these MICOS subunits are essential (not dispensable) for proper crista formation

4) Surprisingly, neither MICOS subunit is essential for in vitro growth or differentiation from ABS to sexual stages, and from the latter to sporozoites. This says more about the biology of plasmodium itself than anything about the essentiality of Mic60, ie plasmodium life cycle progression tolerates defects to mitochondrial morphology. But yes, I agree with the authors that Mic60's apparent insignificance for cell growth in examined conditions does differ with its essentiality in other eukaryotes. But fitness costs were not assayed (eg by competition between mutants and WT in infection of mosquitoes)

5) Decreased fitness of the mutants is implied by a reduction of oocyte formation. 

While interesting in their own way, collectively they do not represent a major advance in our understanding of MICOS evolution. Furthermore, the findings bifurcate into categories informing MICOS or Plasmodium biology. Both aspects are somewhat underdeveloped in their current form. 

This is unfortunate because there seem to be many missed opportunities in the manuscript that could, with additional experiments, lead to a manuscript with much wider impact. For me, what is remarkable about Plasmodium MICOS that sets it apart from other iterations is the apparent absence of the Mic10 subunit. Purification of plasmodium MICOS via the epitope tagged Mic60 and Mic19 could have verified that MICOS is assembled without this core subunit. Perhaps Mic60 and Mic19 are the vestiges of the complex, and thus operate alone in shaping cristae. Such a reduction may also suggest the declining importance of mitochondria in plasmodium.

Another missed opportunity was to assay the impact of MICOS-depletion of OXPHOS in plasmodium. This is a salient issue as maybe crista morphology is decoupled from OXPHOS capacity in Plasmodium, which links to the apparent tolerance of mitochondrial morphology in cell growth and differentiation. I suggested in section A experiments to address this deficit. 

Finally, the authors could assay fitness costs of MICOS-ablation and associated phenotypes by assaying whether mosquito infectivity is reduced in the mutants when they are directly competing with WT plasmodium. Like the authors, I am also surprised that MICOS mutants can pass population bottlenecks represented by differentiation events. Perhaps the apparent robustness of differentiation may contribute plasmodium's remarkable ability to adapt. 

I realize that the authors put a lot of efforts into their study and again, I am very impressed by the sophistication of the methods employed. Nevertheless, I think there is still better ways to increase the impact of the study aside from overinterpreting the conclusions from the data. But this would require more experiments along the lines I suggest in Section A and here.

We thank the reviewer for their extensive analysis of the significance of our findings, including the compliments on our microscopy images and the sophisticated experimental approaches. We hope we have convincingly argued why we could or could not include some of the additional analyses suggested by the reviewer in section 1 above.

With regard to the significance statement, we want to point out that our finding that PfMICOS is not needed for initial formation of cristae (as opposed to organization thereof), is a confirmation of something that has been assumed by the field, without being the actual focus of studies. We argue that the distinction between formation and organization of cristae is important and deserves some attention within the manuscript. The result of MICOS not being involved in the initial formation of cristae, we argue to be relevant in Plasmodium biology and beyond. As for the insights into how MICOS works in Plasmodium we have confirmed that the previously annotated PfMIC60 is indeed involved in the organization of cristae. Furthermore, we have identified and characterized PfMIC19. These findings, we argue, are indeed meaningful insights into PfMICOS.

Reviewer #3 (Evidence, reproducibility and clarity (Required)): 

Summary: 

MICOS is a conserved mitochondrial protein complex responsible for organising the mitochondrial inner membrane and the maintenance of cristae junctions. This study sheds first light on the role of two MICOS subunits (Mic60 and the newly annotated Mic19) in the malaria parasite Plasmodium falciparum, which forms cristae de novo during sexual development, as demonstrated by EM of thin section and electron tomography. By generating knockout lines (including a double knockout), the authors demonstrate that knockout of both MICOS subunits leads to defects in cristae morphology and a partial loss of cristae junctions. With a formidable set of parasitological assays, the authors show that despite the metabolically important role of mitochondria for gametocytes, the knockout lines can progress through the life stages and form sporozoites, albeit with diminished infection efficiency. 

We thank the reviewer for their time and compliment.

Major comments: 

1) The authors should improve to present their findings in the right context, in particular by:

(i) giving a clearer description in the introduction of what is already known about the role of MICOS. This starts in the introduction, where one main finding is missing: loss of MICOS leads to loss of cristae junctions and the detachment of cristae membranes, which are nevertheless formed, but become membrane vesicles. This needs to be clearly stated in the introduction to allow the reader to understand the consistency of the authors' findings in P. falciparum with previous reports in the literature.

We extended the introduction to include this information.

(ii) at the end to the introduction, the motivating hypothesis is formulated ad hoc "conclusive evidence about its involvement in the initial formation of cristae is still lacking" (line 83). If there is evidence in the literature that MICOS is strictly required for cristae formation in any organism, then this should be explained, because the bona fide role of MICOS is maintenance of cristae junctions (the hypothesis is still plausible and its testing important).

To clarify we rephrased the sentence to: “Although MICOS has been described as an organizer of crista junctions, its role during the initial formation of nascent cristae has not been investigated.”

2) Line 96-97: "Interestingly, PfMIC60 is much larger than the human MICOS counterpart, with a large, poorly predicted N-terminal extension." This statement is lacking a reference and presumably refers to annotated ORFs. The authors should clarify if the true N-terminus is definitely known - a 120kDa size is shown for the P. falciparum but this is not compared to the expected length or the size in S. cerevisiae.
To solve the reference issue, we added the uniprot IDs we compared to see that the annotated ORF is bigger in Plasmodium. We also changed the comparison to yeast instead of human, because we realized it is confusing to compare to yeast all throughout the figure, but then talk about human in this specific sentence.

Regarding whether the true N-terminus is known. Short answer: No, not exactly.

However, we do know that the Pf version is about double the size of the yeast protein.

As the reviewer correctly states, we show the size of 120kDa for the tagged protein in Figure 1G. Considering that we tagged the protein C-terminally, and observed a 120kDa product on western blot, it is safe to conclude that the true N-terminus does not deviate massively from the annotated ORF, and hence, that there is a considerable extension of the protein beyond a 60kDa protein. We do not directly compare to yeast MIC60 on our western blots, however, that comparison can be drawn from literature: Tarasenko et al., 2017 showed that purified MIC60 running at ~60kDa on SDS-PAGE actively bends membranes, suggesting that in its active form, the monomer of yeast MIC60 is indeed 60kDa in size.

To clarify, we now emphasize that we ran the Alphafold prediction on the annotated open reading frame (annotated and sequenced by Bohme et al. and Chapell et al. now cited in the manuscript), and revised the wording to make clear what we are comparing in which sentence.

3) lines 244-245: "Furthermore, our data indicates the effect size increases with simultaneous ablation of both proteins?". The authors should explain which data they are referring to, as some of the data in Fig 3 and 4 look similar and all significance tests relate to the wild type, not between the different mutants, so it is not clear if any overserved differences are significant. The authors repeat this claim in the discussion in lines 368-369 without referring to a specific significance test. This needs to be clarified.

As a reply to this and other comments from the reviewers we added the multiple testing within all samples. In addition, to clarify statistics used we included a supplementary dataset with all p-values and statistical tests used.

4) lines 304-306: "Though well established as the cristae organizing system, the role of MICOS in initial formation of cristae remains hidden in model organisms that constitutively display cristae.". This sentence is misleading since even in organisms that display numerous cristae throughout their life cycle, new cristae are being formed as the cells proliferate. Thus, failure to produce cristae in MICOS knockout lines would have been observable but has apparently not been reported in the literature. Thus, the concerted process in P. falciparum makes it a great model organism, but not fundamentally different to what has been studied before in other organisms.

We deleted this statement.

5) lines 373-378. "where ablation of just MIC60 is sufficient to deplete functionality of the entire MICOS (11, 15),". The authors' claim appears to be contrary to what is actually stated in ref 15, which they cite:

"MICOS subunits have non-redundant functions as the absence of both MICOS subcomplexes results in more severe morphological and respiratory growth defects than deletion of single MICOS subunits or subcomplexes."

This seems in line with what the authors show, rather than "different".

This sentence has been removed.

6) lines 380-385: "... thus suggesting that membrane invaginations still arise, but are not properly arranged in these knockout lines. This suggests that MICOS either isn't fully depleted,...". These conclusions are incompatible with findings from ref. 15, which the authors cite. In that study, the authors generated a ∆MICOS line which still forms membrane invaginations, showing that MICOS is not required at all for this process in yeast. Hence the authors' implication that MICOS needs to be fully depleted before membrane invaginations cease to occur is not supported by the literature.

This sentence has been deleted in the revised version of the manuscript.

Minor comments:

7) The authors should consider if the first part of their title could be seen as misleading: It suggests that MICOS is "the architect" in cristae formation, but this is not consistent with the literature nor their own findings.

Title is changed accordingly

Minor comments:

- Line 43, of the three seminal papers describing the discovery of MICOS in 2011, the authors only cite two (refs 6 and 7), but miss the third paper, Hoppins et al, PMID: 21987634, which should probably be corrected.

Done, the paper is now cited

- Page 2, line 58: for a more complete picture the authors should also cite the work of others here which shows that although at very low levels, e.g. complex III (a drug target) and ATP synthase do assemble (Nina et al, 2011, JBC).

Done

- Page 3, line 80: "Irrespective of the shape of an organism's cristae, the crista junctions have been described as tubular channels that connect the cristae membrane to the inner boundary membrane (22, 24)." This omits the slit-shaped cristae junctions found in yeast (Davies et al, 2011, PNAS), which the authors should include.

The paper and concept have been added to the manuscript, though the sentence has been moved up in the introduction, when crista junctions are first introduced.

- Line 97: "poorly predicted N-terminal extension", as there is no experimental structure, we don't know if the prediction is poor. Presumably the authors mean either poorly ordered or the absence of secondary structure elements, or the poor confidence score for that region in the prediction? This should be clarified or corrected.

We were referring to the poor confidence score. To address this comment as well as major point 2, we rewrote the respective paragraph. It now clearly states that confidence of the prediction is low, and we mention the tool that was used to identify conserved domains (Topology-based Evolutionary Domains).

- Line 98: "an antiparallel array of ten β-sheets". They are actually two parallel beta-sheets stacked together. The authors could find out the name of this fold, but the confidence of the prediction is marked a low/very low. So, its existence is unknown, not just its "function".

We adapted the domain description to “a stack of two parallel beta-sheets" and replaced the statement on unknown function by the statement “Because this domain is predicted solely from computational analysis, both its actual existence in the native protein and its biological function remain unknown.”

- Fig 1B: The authors show two alphafold predictions of S. cerevisiae and P. falciparum Mic60 structures. There is however an experimental Mic60/19 (fragment) structure from the former organism (PMID: 36044574), which should be included if possible
We appreciate the reviewer’s suggestion and note that the available structural data indeed provides valuable insight into how MIC60 and MIC19 interact. However, these structures represent fusion constructs of limited protein fragments and therefore capture only a small portion of each protein, specifically the interaction interface. Because our aim in Fig. 1B is to compare the overall domain architecture of the full‑length proteins, we believe that including fragment‑based structures would be less informative in this context.

- Line: 318-321: "The same trend was observed for PfMIC19 and PfMIC60. Although transcriptomic data suggested that low-level transcripts of PfMIC19 and PfMIC60 are present in ABS (38), we did not detect either of the proteins in ABS by western blot analysis. While this statement is true, the authors should comment on the sensitivity of the respective methods - how well was the antibody working in their hands and how do they interpret the absence of a WB band compared to transcriptomics data? 
The HA antibody used in our experiments is a standard commercial reagent that performs reliably in both WB and IFA, although it shows a low background signal in gametocytes. We agree that the sensitivity of the method and the interpretation of weak or absent bands should be addressed explicitly. Transcript levels for both PfMIC19 and PfMIC60 in asexual blood stages fall within the <25 percentile, suggesting that these signals likely represent background. Nevertheless, we acknowledge that low‑level protein expression below the detection limit of western blot analysis cannot be excluded. To reflect these considerations, we added the sentence: ‘The apparent absence could indicate that transcripts are not translated in ABS or that the proteins’ expression was below detection limits of western blot analysis.

- Lines 322-323: would the authors not typically have expected an IFA signal given the strength of the band in Western blot? If possible, the authors should comment if the negative fluorescence outcome can indeed be explained with the low abundance or if technical challenges are an equally good explanation.

Considering the nature of the investigated proteins (embedded in the IMM and spread throughout the mitochondria) difficulties in achieving a clear signal in IFA or U-ExM are not very surprizing. While epitopes may remain buried in IFA, U-ExM usually increases accessibility for the antibodies. However, U-ExM comes at the cost of being prone to dotty background signals, therefore potentially hiding low abundance, naturally dotty signals such as the signal of MICOS proteins that localize to distinct foci (at the CJ) along the mitochondrion. Current literature suggests that, in both human and yeast, STED is the preferred method for accurate spatial resolution of MICOS proteins (<https://www.ncbi.nlm.nih.gov/pubmed/32567732>,<https://www.ncbi.nlm.nih.gov/pubmed/32067344>). Unfortunately, we do not have experience with, nor access to, this particular technique/method.

- Lines 357-365: the authors describe limitations of the applied methods adequately. Perhaps it would be helpful to make a similar statement about the analysis of 3D objects like mitochondria and cristae from 2D sections. E.g. the apparent cristae length depends on whether cristae are straight (e.g. coiled structures do not display long cross sections despite their true length in 3D).
The limitations of other methods are described in the respective results section.

We added a clarifying sentence in the results section of Figure 4:

“Note that such measurements do not indicate the true total length or width of cristae, as the data is two-dimensional. The recorded values are to be considered indicative of possible trends, rather than absolute dimensions of cristae.“

This statement refers to the length/width measurements of cristae.

In the context of Figure 4 D we mention the following (see preprint lines 229 – 230): “We expect this effect to translate into the third dimension and thus conclude that the mean crista volume increases with the loss of either *PfMIC19*, *PfMIC60*, or both.”

For Figure 5, we included a clarifying statement in the results section of the preprint (lines 269 – 273): “Note that these mitochondrial volumes are not full mitochondria, but large segments thereof. As a result of the incompleteness of the mitochondria within the section, and the tomography specific artefact of the missing wedge, we were unable to confirm whether cristae were in fact fully detached from the boundary membrane, or just too long to fit within the observable z-range. “

- Line 404: perhaps undetected or similar would be a better description than "hidden"?
 The sentence does not exist in the revised manuscript

Reviewer #3 (Significance (Required)):

The main strength of the study is that it provides the first characterisation of the MICOS complex in P. falciparum, a human parasite in which the mitochondrion has been shown to be a drug target. Mic60 and the newly annotated Mic19 are confirmed to be essential for proper cristae formation and morphology, as well as overall mitochondrial morphology. Furthermore, the mutant lines are characterised for their ability to complete the parasite life cycle and defects in infection effectivity are observed. This work is an important first step for deciphering the role of MICOS in the malaria parasite and the composition and function of this complex in this organism. The limitation of the study stems from what is already known about MICOS and its subunits in

great detail in yeast and humans with similar findings regarding loss of cristae and cristae defects. The findings of this study do not provide dramatic new insight on MICOS function or go substantially beyond the vast existing literature in terms of the extent of the study, which focuses on parasitological assays and morphological analysis. Exploring the role of MICOS in an early-divergent organism and human parasite is however important given the divergence found in mitochondrial biology and P. falciparum is a uniquely suited model system. One aspect that would increase the impact of the paper would be if the authors could mechanistically link the observed morphological defects to the decreased infection efficiency, e.g. by probing effects on mitochondrial function. This will likely be challenging as the morphological defects are diverse and the fitness defects appear moderate/mild.

As suggested by Reviewer 2, we examined mitochondrial membrane potential in gametocytes using MitoTracker staining and did not observe any obvious differences associated with the morphological defects. At present, additional assays to probe mitochondrial function in P. falciparum gametocytes are not sufficiently established, and developing and validating such methods would require substantial work before they could be applied to our mutant lines. For these reasons, a more detailed mechanistic link between the observed morphological changes and the reduced infection efficiency is currently beyond reach.

The advance presented in this study is to pioneer the study of MICOS in P. falciparum, thus widening our understanding of the role of this complex to different model organism. This study will likely be mainly of interest for specialised audiences such as basic research parasitologists and mitochondrial biologists. My own field of expertise is mitochondrial biology and structural biology.
