## Supplementary Figures for "The role of MICOS in organizing mitochondrial cristae in malaria parasites"

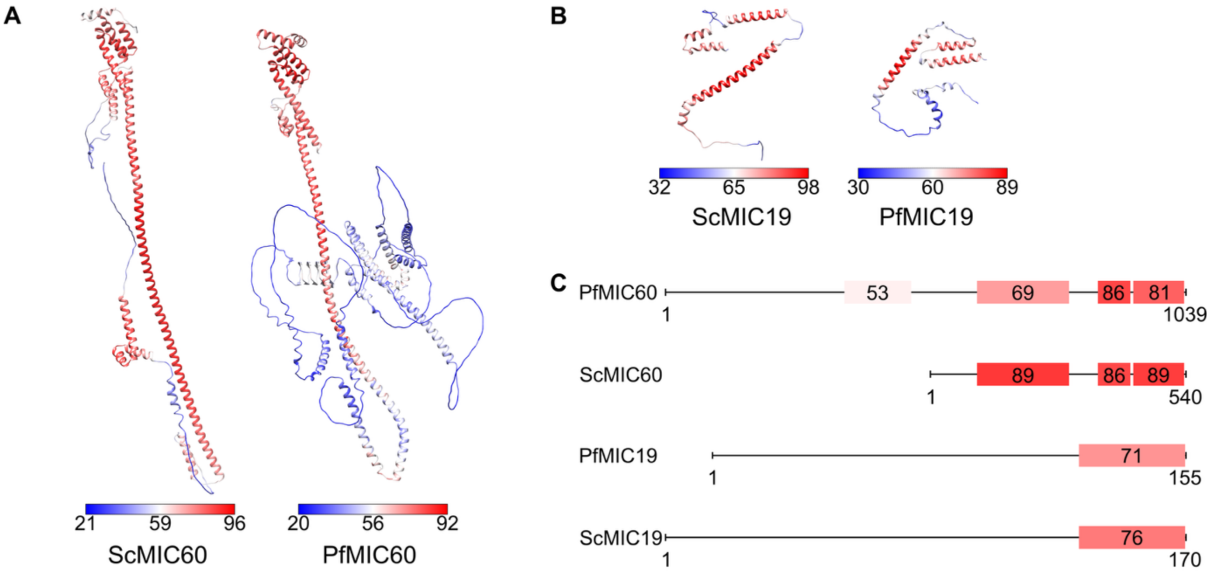


***Figure S1. Overview of local pLDDT confidence scores of the AlphaFold structures shown for MIC60 and MIC19 (Figure 1).*** *A) AlphaFold predictions of ScMIC60 and PfMIC60 (retrieved from the AlphaFold Protein Structure Database entries AF-P36112-F1-v4 and AF-Q8IJW5-F1-v4) (27, 28). Colour scheme follows the pLDDT score from blue (low confidence) to red (high confidence). Total pLDDT range for each model is annotated below the respective prediction. B) AlphaFold predictions of ScMIC19 and PfMIC19 (retrieved from the AlphaFold Protein Structure Database entries AF-P43594-F1-v4 and AF-Q8IIR5-F1-v4) (27, 28). Colour scheme follows the pLDDT score from blue (low confidence) to red (high confidence). Total pLDDT range for each model is annotated below the respective prediction. C) Schematic overview of the predicted domain architectures for A and B. Colour code of the annotated domains follows a global colour scheme from 0 to 100 is represented in blue (theoretical pLDDT of 0) to red (theoretical pLDDT of 100). Exact pLDDT scores of the individual domains are given within the respective box.*

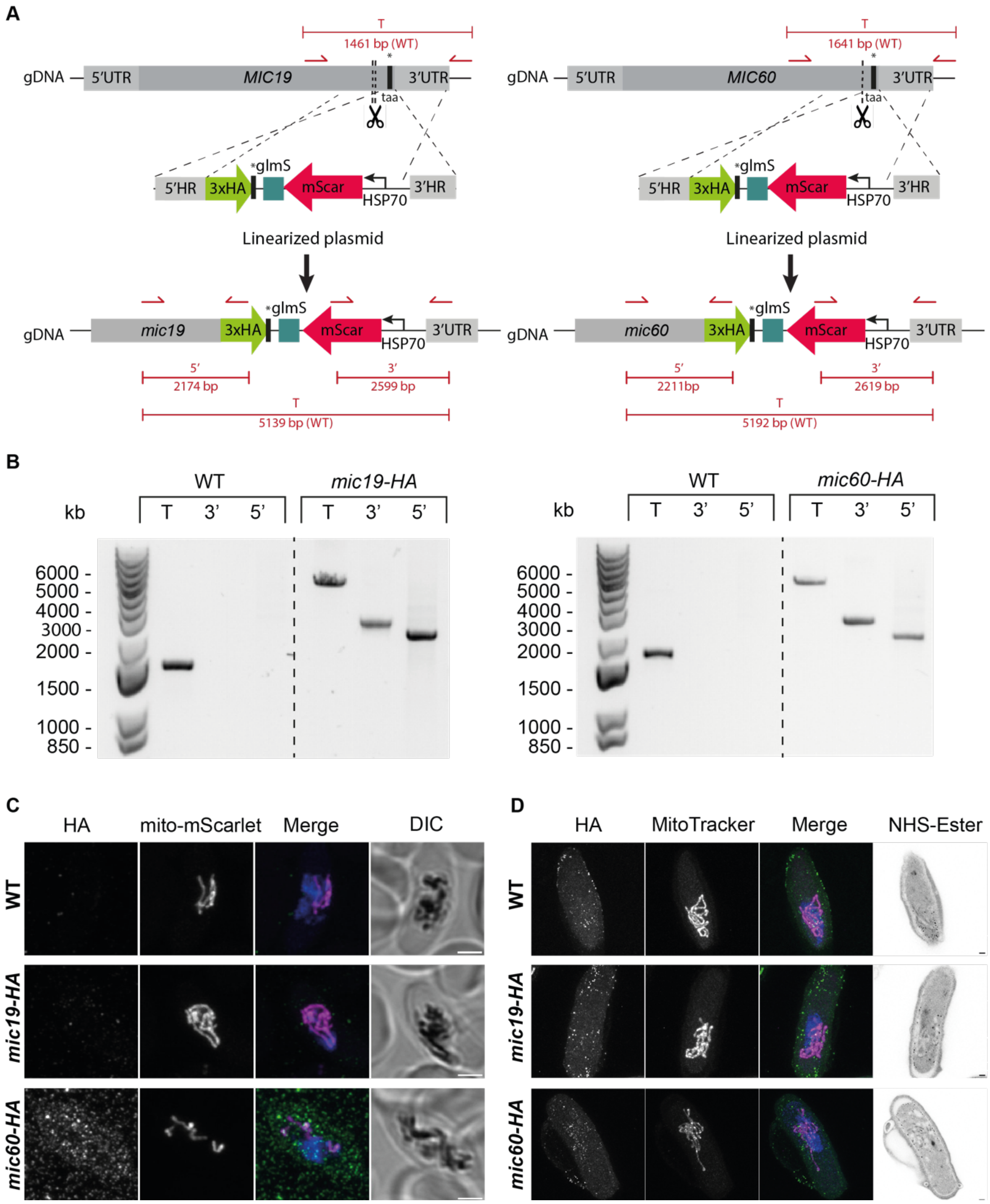


***Figure S2. Generation and verification of mic19-HA and mic60-HA parasite lines and fluorescence microscopy to co-localize PfMIC19 and PfMIC60 with mitochondria.*** *A) Schematic overview of the strategy to generate a parasite line presenting an 3xHA tag on MIC19 or MIC60. CRISPR-Cas9 was used for this approach to generate double-strand breaks (two for MIC19 and one for MIC60; indicated by scissors). The repair plasmid contains a 3xHA followed by a GlmS and a bidirectional 3'UTR (not shown in figure, PBANKA_142660). The guide sequence was recodonised to prevent the guides from recognizing the target after integration. In addition, the construct presents an mScarlet fused with the targeting sequence of the mitochondrial protein HSP70-3 (PF3D7_1134000) under the HSP70-3*

*promoter. B) Diagnostic PCR of mic19-HA and mic60-HA parasite lines with WT control and integration-specific primers (indicated in red in panel A). C) Fluorescence microscopy of fixed WT (MitoRed) mic19-HA and mic60-HA, stained with rat anti-HA (green) and DAPI for DNA (blue). Mitochondria are visible due to the presence of mitochondrial-mScarlet (magenta). Images are representative of the observation during the experiments. All images are a maximum projection of a Z-stack. DIC shows a slice in the middle of the Z-stack to show parasite shape. Scale bar = 2 $\mu$m. D) U-ExM of WT (MitoRed), mic19-HA and mic60-HA, stained with rat anti-HA with biotin amplification (green), MitoTracker to visualize mitochondria (magenta), SYTOX for the DNA (blue) and NHS-ester for amine reactive groups (shades of grey). Scale bar = 2 $\mu$m.*

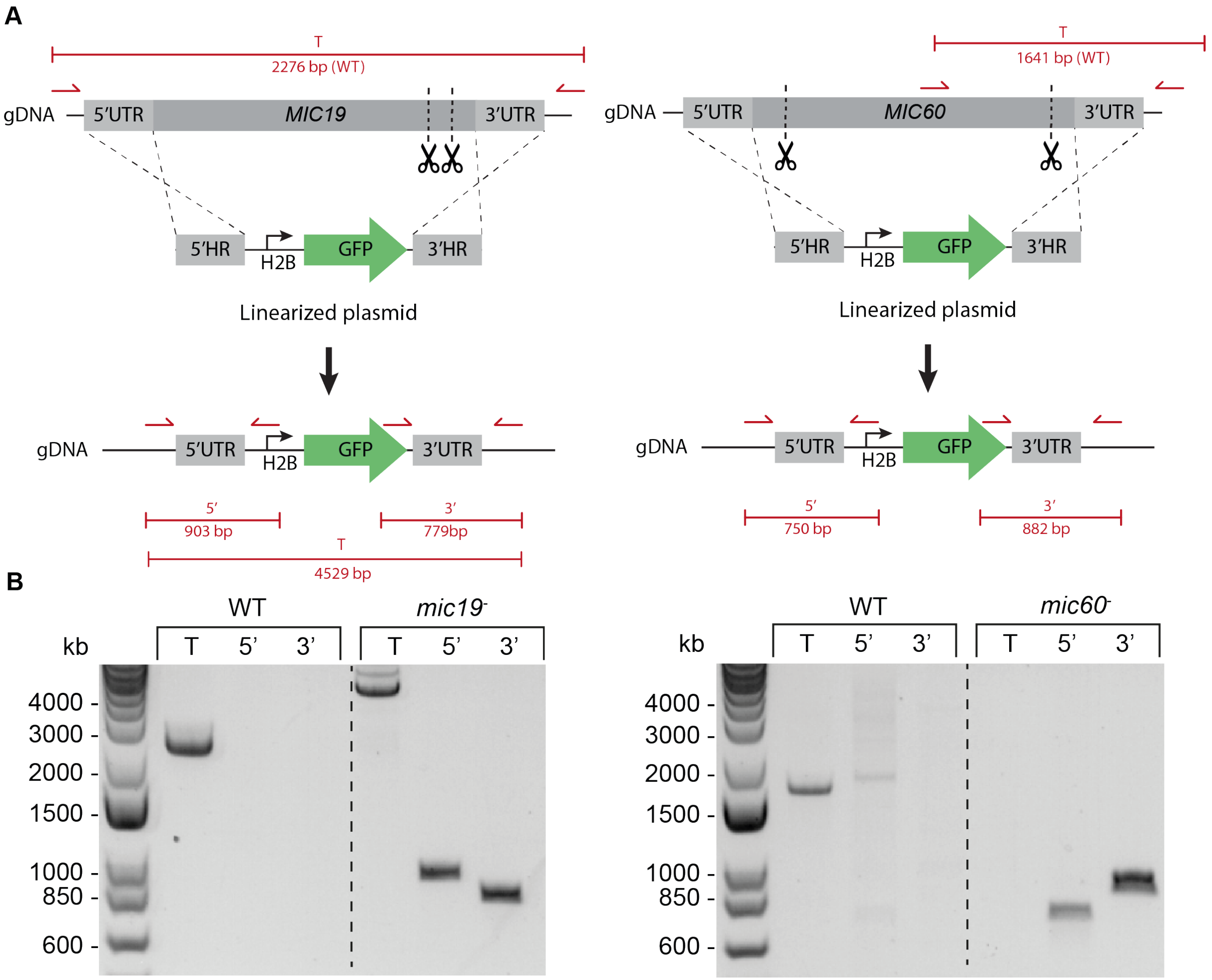


***Figure S3. Generation and verification of mic19⁻ and mic60⁻ parasite lines**. A) Schematic overview of the strategy to generate a knockout parasite line for MIC19 and MIC60. CRISPR-Cas9 was used for this approach to generate two double-strand breaks (indicated by scissors). The repair plasmid contains a GFP under the control of the H2B promoter (PF3D7_1105100) followed by a bidirectional 3'UTR (not shown in figure, PBANKA_142660). B) Diagnostic PCR of mic19 ⁻ and mic60 ⁻ parasite lines with WT (NF54) control and integration- specific primers (indicated in red in panel A).*

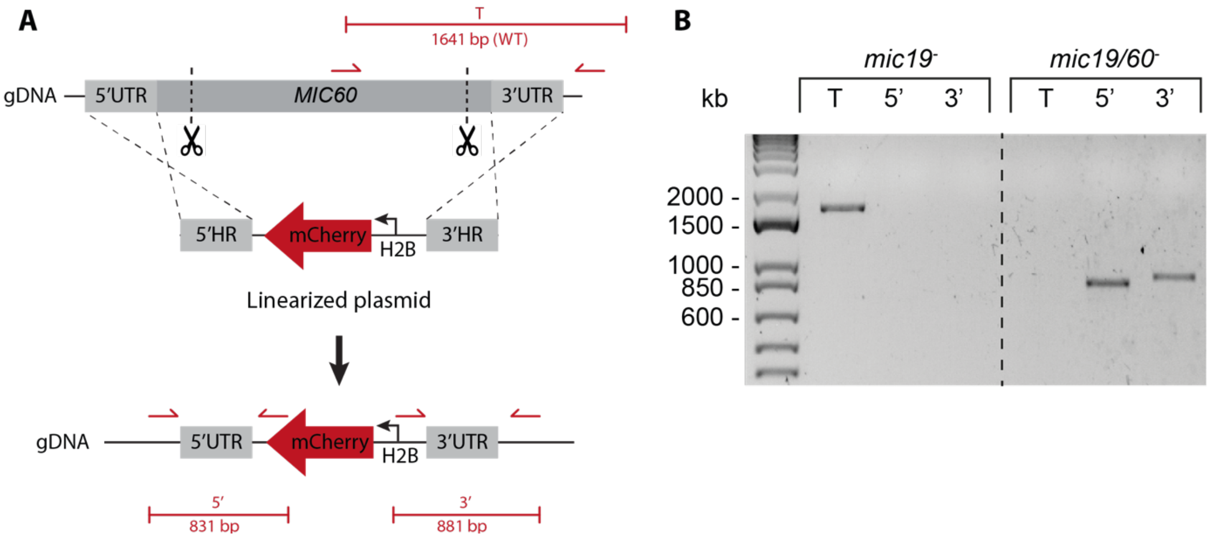


***Figure S4. Generation and verification of mic19/60⁻ parasite line****.* *A) Schematic overview of the strategy to generate the double knockout parasite line with mic19⁻ as genetic background. CRISPR-Cas9 was used for this approach to generate two double-strand breaks (indicated by scissors). The repair plasmid contains an mCherry under the control of the H2B promoter (PF3D7_1105100) followed by a bidirectional UTR (not shown in figure, PBANKA_142660). B) Diagnostic PCR of mic19/60⁻ parasite lines with mic19⁻ control and integration-specific primers (indicated in red in panel A).*

**A**

| **Exp n°** | **NF54** | | ***mic19⁻*** | | ***mic60⁻*** | | ***mic19/60⁻*** | |
|---|---|---|---|---|---|---|---|---|
| | % inf mosq. | mean | % inf mosq. | mean | % inf mosq. | mean | % inf mosq. | mean |
| Exp1 | 100% (20/20) | 120,3 | 100% (20/20) | 8,9 | | | | |
| Exp2 | 100% (20/20) | 16,1 | 95% (19/20) | 13,2 | | | | |
| Exp3 | 100% (20/20) | 27,3 | 85% (17/20) | 5,8 | 95% (19/20) | 13,2 | 95% (19/20) | 25,2 |
| Exp4 | 100% (10/10) | 63,9 | | | 90% (9/10) | 37,6 | | |
| Exp5 | 95% (19/20) | 5,7 | | | 80% (16/20) | 2,25 | 60% (12/20) | 2,2 |
| Exp6 | 100% (10/10) | 68 | | | | | 100% (10/10) | 18 |

**B**

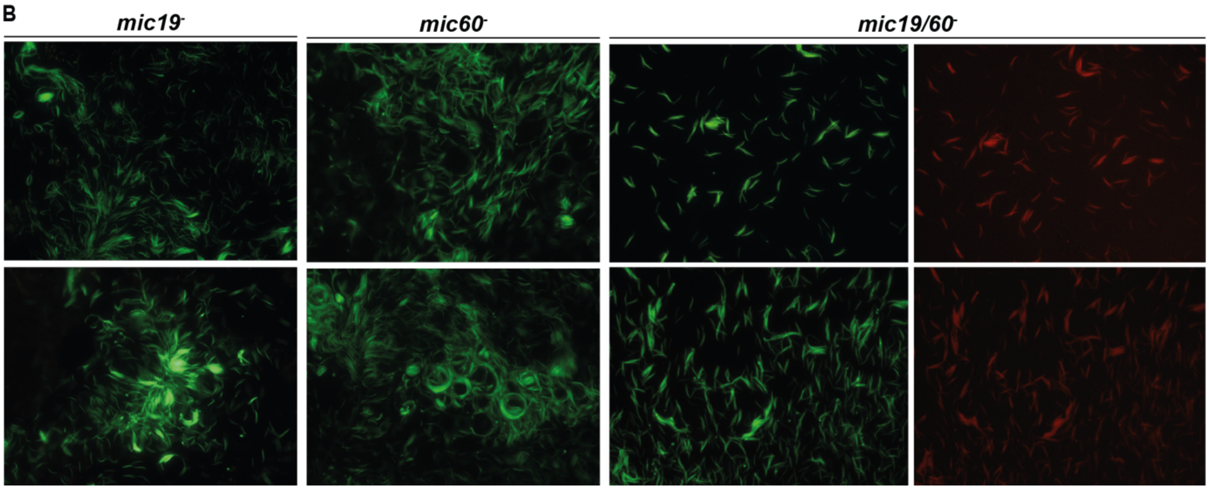


***Figure S5. Live microscopy pictures of mic19⁻, mic60⁻ or mic19/60⁻ sporozoites.*** *A). Summary of percentage of infected mosquitoes, n° of mosquito counted and mean oocyst count per midgut in every independent experiment displayed in Figure 2C. B) Images of mic19⁻, mic60⁻, and mic19/60⁻ sporozoites acquired with a Zeiss Axioscope with AxioCam iCc1 camera and a 40X or 100X objective using the Zen Blue (version 2.5.75.0) software and processed using Fiji with identical settings.*

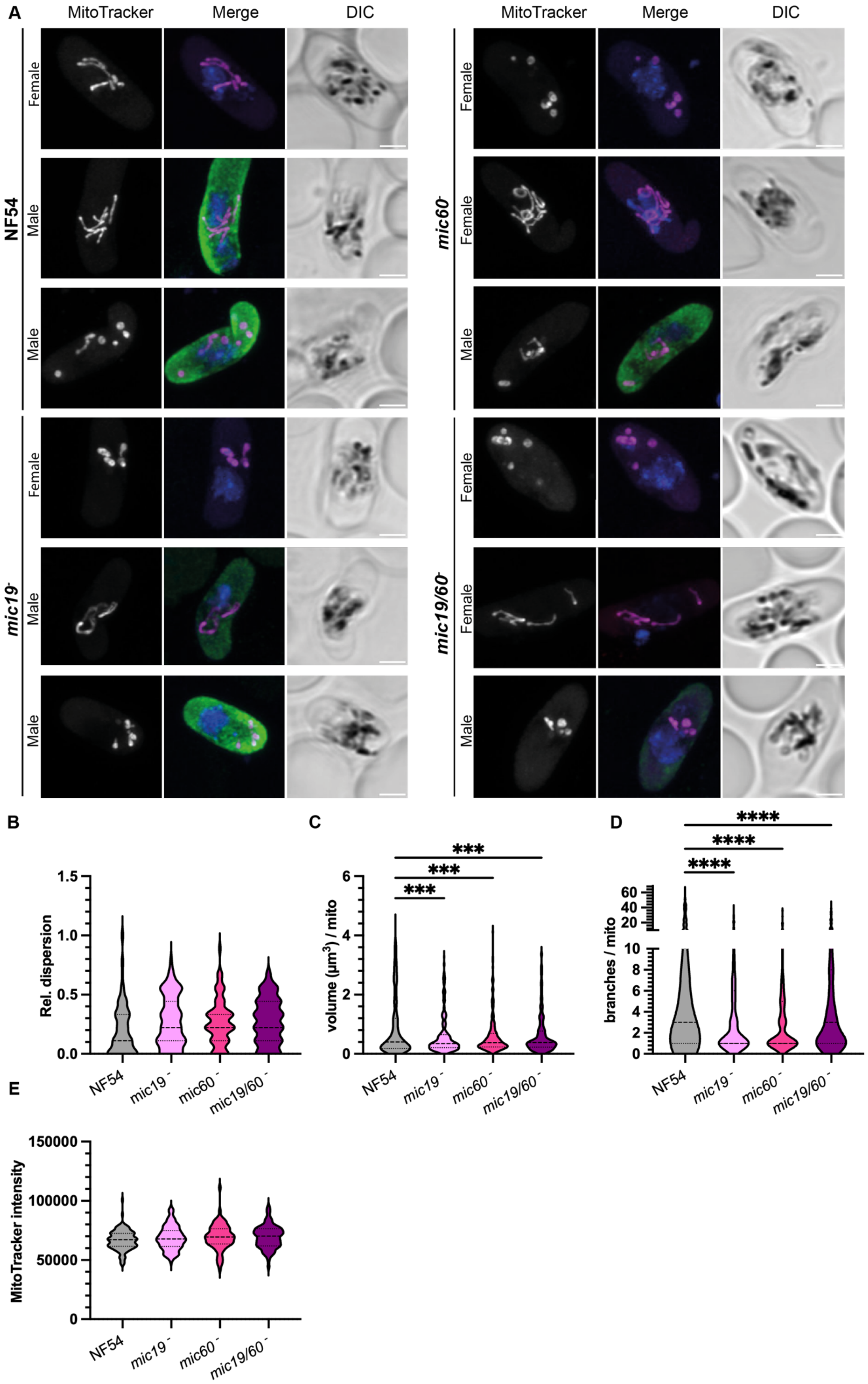

A
MitoTracker
Merge
DIC
NF54
Female
Male
Male
mic60-
mic19-
mic19/60-
B
Rel. dispersion
1.5
1.0
0.5
0.0
NF54
mic19 -
mic60 -
mic19/60 -
C
volume (µm³) / mito
6
4
2
0
***
***
***
D
branches / mito
60
40
20
10
8
6
4
2
0
****
****
****
E
MitoTracker intensity
150000
100000
50000
0

***Figure S6. Fluorescence microscopy of mature* P. falciparum *gametocytes of mic19⁻, mic60⁻ and mic19/60⁻ shows aberrant mitochondrial morphology.*** *A) Representative fluorescence microscopy images of mature parental (NF54), mic19⁻, mic60⁻, and mic19/60⁻ gametocytes. Samples were stained with MitoTracker (magenta) for the mitochondria, DAPI (blue) for DNA and alpha tubulin II (green) as a male marker. Images are maximum intensity projections of Z-stack confocal Airyscan images. DIC shows a single slice of the Z-stack. Scale bars: 2 µm. (B-F) Quantification of (B) relative mitochondrial dispersion calculated as (n° of detected mitochondrial signals -1) / (n° of maximum detected mitochondrial signals -1). Values range from 0 (no dispersion) to 1 (maximal dispersion). (D) volume per mitochondrial signal, (E) branches per mitochondrion (log), (F) MitoTracker signal intensity in the same samples as those used in Figure 3 to estimate differences in membrane potential. In all cases n° of replicates > 3 with total number of cells ≥ 75. Statistical significance calculated with one-way ANOVA with Sidàk correction for full comparison between all cell lines; significance is indicated as follows: *p < 0.05, **p < 0.01, ***p < 0.001, ****p < 0.0001.*

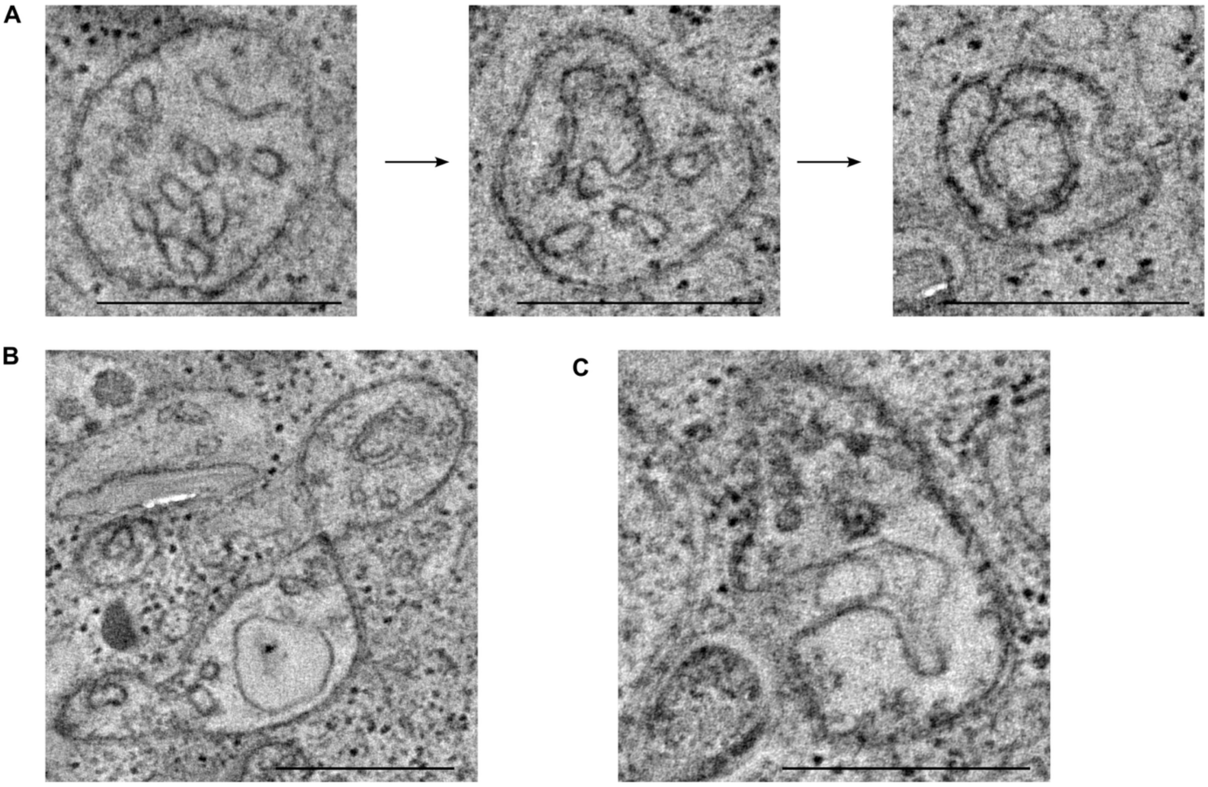


***Figure S7.* P. falciparum *gametocytes of the mic19/60⁻ line show a variety of aberrant cristae phenotypes.*** *A) Single TEM tomography slices of three different mitochondria with aberrant cristae. Depicted from left to right are examples of cristae of increasing length. While the mitochondrion on the left shows a long and slender crista, the longest crista visible in the central panel is not only long but almost closes to a ring shape. The micrograph on the right shows a fully closed, double-membrane ring with a connecting crista junction to the boundary membrane. B – C) Examples of aberrant cristae phenotypes as observed in TEM tomography. Scale bars: 500 nm.*

***Table S1. Primer and guide RNA sequences for integration PCR and the generation of repair and guide plasmids****. Abbreviation used: HR = homology region, F = forward, R = reverse. In red represented the overhang for restriction sites and in green the gRNA sequences. "/" indicates that the primer has been used for different purposes.*

| Primer name | Primer function | Sequence | Restriction site |
|---|---|---|---|
| Generation of repair plasmids | | | |
| ST_54 | 5' HR *mic19-HA* F | tttttCTCGAGATGGGTAACACATTTTCATCTTATGACGAAAAAGAGAG | XhoI |
| ST_55 | 5' HR *mic19-HA* R | ttttttaGGATCCACCTGATCCGCCGATTTTAATTCTTTGTACACAACTTTCATATTCTCTTAAATATTTATAACATCTGGAATATGAATTTAAAAATCCAAAAGTAG | BamHI |
| ST_56 | 3' HR *mic19-HA*/- F | aaaaaaGAATTCTAAAAAATATATTCTGTTCATAAGAATC | EcoRI |
| ST_57 | 3' HR *mic19-HA*/- R | ttttttGCCGGCAGATTGTACTTATTAAGGAGCG | NgoMIV |
| ST_60 | 5' HR *mic60-HA* F | tatataCTCGAGGAATTAGAAAAAGAGAAAAACAAAATTCAACAAC | XhoI |
| ST_61 | 5' HR *mic60-HA* R | tatataGGATCCACCTGATCCGCCCTCACACAATGTCAATGTTTTACTAGCAAGCATTATTCTAGAAACAACTAACCTAAGATAAAATTTAAACAAGATAAC | BamHI |
| ST_62 | 3' HR *mic60-HA*/- F | tattaaGAATTCCAAATTTACAAACTATTATATTTACACATCACAATC | EcoRI |
| ST_63 | 3' HR *mic60-HA*/- R | aaaaaaGCCGGCGGGAAGAAATAAAAATACTGCATGCG | NgoMIV |
| ST_75 | 5' HR *mic19*$^-$ F | aaaaaaCCCGGGGAAAAATGCATATATTAAAATGATCAATGGAATATAAG | XmaI |
| ST_74 | 5' HR *mic19*$^-$ R | atatatCTCGAGCGTATCCTTTTGAAAAATTCAAAAAGGG | XhoI |
| ST_76 | 5' HR *mic60*$^-$ F | atatatCCCGGGCATTTAATCACAAAAGGTACGTTC | XmaI |
| ST_77 | 5' HR *mic60*$^-$ R | ttttttCTCGAGGATTAATGTTAATATTTCCTTTGCTC | XhoI |
| Guide RNA sequences | | | |
| ST_58 | MIC19 guide 1 F | TATTGCCTCAAATATTTATAACACC | |
| ST_59 | MIC19 guide 1 R | AAACGGTGTTATAAATATTTGAGGC | |
| ST_44 | MIC19 guide 2 F | TATTGCCAGGTGTTATAAATATTTG | |
| ST_45 | MIC19 guide 2 R | AAACCAAATATTTATAACACCTGGC | |
| ST_50 | MIC60 guide 1 F | TATTGATCTTAGGTTAGTTGTCTCA | |
| ST_51 | MIC60 guide 1 R | AAACTGAGACAACTAACCTAAGATC | |
| ST_52 | MIC60 guide 2 F | TATTGAGATGGTTTACGAGATGTCG | |
| ST_53 | MIC60 guide 2 R | AAACCGACATCTCGTAAACCATCTC | |
| Diagnostic Integration PCR | | | |
| ST_68 | 5' *mic60-HA* F | GAGAAAGAAAAAGAAAAGTTTC | |
| ST_70 | 5' *mic19-HA* F | AATTTTTCAAAAGGATACGC | |
| JV_64 | 5' *mic19-HA* and *mic60-HA* R | ATGGTGAGCAAGGGCGAGG | |
| ST_79 | 5' *mic60*$^-$ F | AAGGAAAAAGGAAAAATATTTATGTAC | |
| ST_78 | 5' *mic19*$^-$ F | TTATATTTGTCCTGATCTTTAAATGG | |
| NP_50 | 5' *mic19*$^-$ and *mic60*$^-$ R | CTTAATATTGATAAGTATCATGTG | |
| ST_69 | 3' *mic60-HA/-* F | TATATTACGCAAAATAAGGG | |
| ST_71 | 3' *mic60-HA/-* F | CTTTTTATTTTTTTAAATGCGC | |
| JV_168 | 3' *mic19-HA* and *mic60-HA* R | TTTAGCTCTACTCTAAAAAACTGATAAAAG | |
| NP_49 | 3' *mic19*$^-$ and *mic60*$^-$ R | CCGAAAAAGTTAAAATTAATTTAC | |
| NP_190 | 3' *mic19/60*$^-$ R | AGTCATATCCAGGAATAAACATAC | |

**Table S2. List of primary and secondary antibodies used during IFA, U-ExM and WB.**

| Antibody | Type | Brand and catalogue number | Dilution |
|---|---|---|---|
| Rat anti-HA | Primary | ThermoFisher, 11867423001 | 1:1000 (WB), 1:250 (U-ExM) |
| Mouse anti-αTubulin | Primary | Invitrogen, Ma1-19162 | 1:500 (IFA) |
| Rabbit anti-HSP70 | Primary | StressMArq Biosciences, SPC-186D | 1:2000 (WB) |
| Donkey Anti-Mouse 647 Alexa fluorophore | Secondary | Invitrogen, A-31571 | 1:500 |
| Biotin, goat anti- rat | Secondary | ThermoFisher,31830 | 1:400 (U-ExM) |
| Streptavidin, Alexa fluorophore 488 | Conjugated | ThermoFisher, 11223 | 1:400 (U-ExM) |
| Goat Anti-Rabbit 680 Fluorophore | Secondary | LICOR, IRDy, 926-68071 | 1:5000 (WB) |
| Goat anti-Rat 800 Fluorophore | Secondary | LICOR, IRDy, 926-32219 | 1:5000 (WB) |
